# Fossil calibrations for molecular analyses and divergence time estimation for true crabs (Decapoda: Brachyura)

**DOI:** 10.1101/2023.04.27.537967

**Authors:** Javier Luque, Heather D. Bracken-Grissom, Javier Ortega-Hernández, Joanna M. Wolfe

## Abstract

True crabs, or Brachyura, comprise over 7,600 known species and are among the most ecologically dominant, economically significant, and popularly recognized group of extant crustaceans. There are over 3,000 fossil brachyuran species known from mid and upper Jurassic, Cretaceous, and Cenozoic deposits across the globe, many of them preserved in exquisite detail, but the origins and early evolution of true crabs remain unresolved. This uncertainty hinders the identification of the stratigraphically earliest occurrence of major brachyuran groups in the fossil record, obscuring our understanding of their phylogenetic relationships and thus the ability to estimate divergence times to answer large scale macroevolutionary questions. Here, we present 36 vetted fossil node calibration points for molecular phylogenetic analysis of crabs (one Anomura and 35 Brachyura) and reassess the earliest occurrences of several key clades based on recent fossil discoveries or re-examination of previous studies. For each calibrated node, we provide the minimum and tip maximum ages for the stratigraphically oldest fossil that can be reliably assigned to the group. Disentangling the anatomical disparity of fossil forms and their phylogenetic relationships is crucial to recognize the earliest branching members among brachyuran groups. This represents a critical first step understanding the evolution of carcinization and decarcinization in true crabs, the appearance of key adaptations, and the transition from sea to land and freshwater. The identification of reliable fossils for deep time calibrations, both as tips and nodes, is pivotal to ensure not only precise but more accurate divergence time estimations when reconstructing the crab tree of life.

**PLAIN LANGUAGE SUMMARY:** We present 36 vetted fossil calibration points for molecular phylogenetic analysis of crabs (one Anomura and 35 Brachyura) and reassess the earliest occurrences of several key groups based on recent fossil discoveries or re-examination of previous studies, together with discussions for each taxon. We also provide some general observations and recommendations on fossil age selection and stratigraphic considerations. The identification of reliable fossils for deep time calibrations, both as tips and nodes, is pivotal to ensure not only precise but more accurate divergence time estimations when reconstructing phylogenetic trees.

## INTRODUCTION

The origins and evolution of true crabs, or Brachyura, have sparked the fascination of scientists and the public alike in the last decades, thanks to their astonishing anatomical diversity (disparity) and multiple convergent instances of gain/loss of the crab-like body plan (i.e., carcinization and decarcinization) (Scholtz, 2014; Luque et al., 2019b; Wolfe et al., 2021). With over 7,600 extant species and more than 3,000 fossil species known (see Ng et al., 2008; Schweitzer et al., 2010; Luque et al., 2019b; Poore and Ahyong, 2023, and references therein), brachyurans are one of the most speciose groups of crustaceans. The monophyly of Brachyura is widely accepted, based on morphological and molecular grounds, and they are considered reciprocal sister groups with Anomura (collectively known as false crabs and allies), both forming a clade referred to as Meiura (Scholtz and Richter, 1995; Schram and Dixon, 2004; Hegna et al., 2020).

The systematics and classification of brachyurans have changed considerably in the last few decades, especially concerning family-level ranks and above. New lines of evidence have become available in the form of comprehensive phylogenetic studies, new fossil discoveries, and the critical re-interpretations of previous ones using novel technologies. Molecular and morphological phylogenetics are key to better understanding the evolutionary relationships through common ancestry among and within brachyurans, allowing us to quantitatively examine previous views and hypotheses largely based on traditional alpha taxonomy. This is particularly important when considering problematic groups such as the so-called podotremes and the heterotreme eubrachyurans, now widely recognized as paraphyletic groups, as well as the relationships among primary freshwater groups (e.g., Tan et al., 2018; Wang et al., 2018; Luque et al., 2019b; Ma et al., 2019; Tang et al., 2020; Wang et al., 2020; Luque et al., 2021; Tsang et al., 2022; Wolfe et al., 2022; Zhang et al., 2022; Poore and Ahyong, 2023).

While molecular and morphological phylogenies provide different types of evidence for understanding evolutionary relationships among organisms, fossils offer critical anatomical, spatial, and temporal information inaccessible from extant species alone (Luque et al., 2019b). Thus, accurate calibrations in a phylogenetic framework require reliable fossils, both in terms of their systematic affinities and their chronostratigraphic occurrences, since the use of equivocal (e.g., superficially similar but unrelated) or poorly constrained temporal and stratigraphic fossils may lead to precise yet inaccurate dating (either as tips or nodes), divergence time estimates, and therefore may confound interpretations of macroevolutionary patterns and processes over time.

The fossil record of brachyurans crabs consists mainly of marine taxa, which tend to be the most abundant and often well preserved. Conversely, fossils of non-marine crabs (i.e., terrestrial, semi-terrestrial, and freshwater) are remarkably scarce and often fragmentary, largely due to the relatively dynamic and high-energy environments they inhabit, the geochemistry of the substrates, and scavenging of their corpses or even re-working and consumption of their own exuviae (e.g., Locatelli, 2013; Luque, 2017; Luque et al., 2018). Such biases limit the number of remains than can fossilize, often restricted to the most resilient biomineralized body parts (e.g., claw dactyli and propodi remains), with whole organisms and soft to lightly-biomineralized tissues only preserved under exceptional conditions (Luque et al., 2019b; Luque et al., 2021). In turn, these biases constrain the number of fossils suitable as reliable calibration points in molecular estimations of time divergence, and consequently the available fossil data that can be used to examine the evolution of terrestrialization and the transition from marine to non-marine habitats (Tsang et al., 2014; Luque et al., 2021; Watson-Zink, 2021; Tsang et al., 2022; Wolfe et al., 2022). Understanding the early origins of marine and non-marine brachyurans by means of their fossil record requires not only new paleontological discoveries, but also a critical re-examination of previous records. This holds true for marine crab fossils as well.

Molecular biologists face limitations when selecting fossils for internal node calibrations related to evaluating specific details of such fossil occurrences, like their chronostratigraphic and lithostratigraphic context, and the reliability of their systematic placement, especially with fragmentary material (Gandolfo et al., 2008; Parham et al., 2012; Wolfe et al., 2016). Among crabs, convergence is ubiquitous, pervasive, and rampant, and there is a trove of extinct and extant groups that superficially may resemble each other, particularly in their dorsal carapaces. This may lead to inappropriate selection of fossils that are not in fact related, and introduce large errors and inaccuracies into the estimation of divergence dates when they are used to calibrate particular nodes and branches.

Fossils collected *in-situ* are often assigned a tentative relative age based on the chronostratigraphic age of the geological formation that contains them. However, geological formations can span tens to hundreds of meters in thickness and hundreds of thousands to millions of years in age, whereas the age of a unitary fossil sample represents a single point in time, making it difficult to constrain its age with respect to the age of the entire lithostratigraphic unit. For example, if a formation is known to be ‘Eocene’ in age, but its base and top have not been better constrained via absolute (radiometric isotope) or relative (e.g., biostratigraphy, stratigraphic position) dating, it means that it could, hypothetically, range chronostratigraphically anywhere from 56.0 to 33.9 Ma. Even if the relative age of such formation is further constrained, for example to ‘lower Eocene (Ypresian)’, the age bracket would range from 56.0 to 47.8 Ma, in which case the fossil would be assigned these as its tip maximum and minimum ages (e.g., several Italian fossil occurrences, see below). This can be circumvented when other components of the fossil assemblages shed light on the biostratigraphic ranges of other micro and/or macrofossils (e.g., the porcellanid crab in calibration 1 here), when reliable radioisotopic data are associated to the same strata as the fossils (e.g., the pseudothelphusid freshwater crab in calibration 16 herein), or if magnetostratigraphic information is available (e.g., the parthenopid crab in calibration 23 herein).

An issue that is often oversimplified, and sometimes hard to circumvent, is the extrapolation of the age range of a formation in a specific locality to other geographically separated localities where the same formation occurs. While, in principle, the horizontality and superposition of beds in a normal sequence would allow for regional correlations, extrapolating ages is not always straightforward. This is particularly important for fossils near the lower and upper boundaries of a lithostratigraphic unit, principally due to the irregular geometry of the basins, unconformities and other erosional surfaces, eustasy, differential subsidence, diachronism, lateral facial changes, and pinching of the strata, to name some. Thus, assigning the age of a formation in one locality to a fossil found in the same formation in a different locality would be better informed by gaining some level of understanding of the local and regional geological and stratigraphic context.

Lastly, unless the ages near the top and/or the bottom of a sequence are known, an absolute age from a sample coming from a bed within the stratigraphic interval can inform about the overall age range of the formation, but does not represent the age of the formation *per se*, and therefore the age of the fossils present in the formation, unless the fossils come from or near the dated bed itself (as in the case in calibration 16 herein). Nonetheless, an approximated age bracket for a fossil occurring between two dated layers is as good of an approximation as possible, given the available data, in which case the minimum and tip maximum ages of the fossil will be informed and constrained by such absolute ages (e.g., as in the pinnotherid crab in calibration 10 herein).

Here we present a set of 36 phylogenetically vetted fossil calibration points for molecular analyses and divergence time estimations of true crabs (one anomuran outgroup, 35 brachyuran ingroups) (based on the crown group topologies inferred by Wolfe et al., 2022) (Fig. 1), and provide information on the type of material investigated, its repository and catalogue number, phylogenetic justification, minimum and tip maximum ages, age justification, and a discussion, following best practices (Parham et al., 2012) (Table 1). For crown Meiura (Anomura + Brachyura) calibrations 1–7, we provide a soft maximum age of 191.8 Ma based on the estimated maximum age for the brachyuran fossil *Eocarcinus praecursor* Withers, 1932, from the Early Jurassic (Pliensbachian) of England (Table 1). Soft maximum age of 100.9 Ma for crown Eubrachyura calibrations 8–19 and 21–36 is based on the estimated maximum age for the crown eubrachyuran fossil *Cretapsara athanata* Luque, *in* Luque et al., 2021, from the mid-Cretaceous (uppermost Albian–lowermost Cenomanian) of Burma (Table 1). Soft maximum age of 97.0 Ma for crown Portunoidea calibration 20 is based on the estimated maximum age for the stem group portunoid fossil *Eogeryon elegius* Ossó, 2016, from the mid-Cretaceous (Cenomanian) of Spain (Luque et al., 2021).

**Figure 1.**
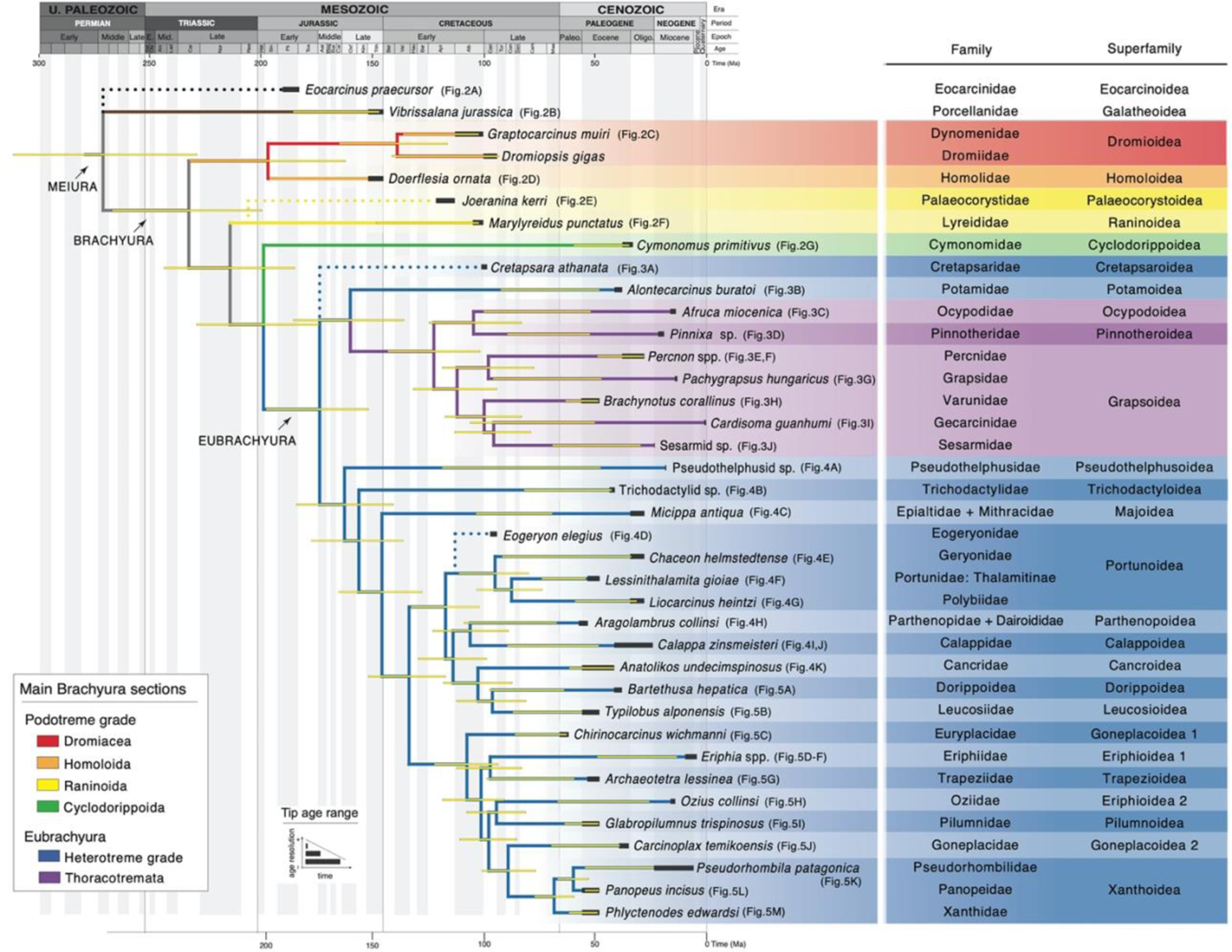
Phylogenetic relationships among 38 meiuran families (36 Brachyura in ingroup), as represented by their oldest reliable vetted fossil species occurrences to date and their age brackets, used as calibration points for node dated molecular divergence time estimations of brachyuran crabs in Wolfe et al. (2022). Fossil species used for node dating are here represented as terminals. Base topology and full maximum clade credibility divergence time estimates (thin shaded yellow bars, representing 95% HPD) after Wolfe et al. (2022). Main brachyuran sections, and the family and superfamily ingroups focus of this study, coloured as follows: Dromiacea (red), Homoloida (orange), Raninoida (yellow), Cyclodorippoida (green), heterotreme Eubrachyura (blue), and thoracotreme Eubrachyura (purple). Oldest crown Porcellanidae (Anomura, brown) and the extinct Eocarcinidae (black dotted line), as non-crown brachyuran outgroups. Fossil age ranges (thick dark grey bars) denote the maximum (left end) and minimum (right end) age brackets for the selected calibration points, based on available chronostratigraphic information derived from isotopic, magnetostratigraphic, or bio-stratigraphic data (see main text). Longer grey bars (e.g., *Calappa zinsmeister*) indicate a wider age gap between the tip maximum and minimum ages for a given taxon, and thus less age resolution for the relative age of the fossil, whereas thinner grey bars (e.g., pseudothelphusid sp.) indicate a narrower age gap between their tip maximum and minimum ages, and thus more resolution and preciseness of the relative age of the fossil. Phylogenetic placement of *Joeranina kerri* (Raninoida: Palaeocorystoidea: Palaeocorystidae) (yellow dotted line) after Luque et al. (2019b) and of *Cretapsara athanata* (Eubrachyura: Cretapsaridae) and *Eogeryon elegius* (Eubrachyura: Portunoidea: Eogeryonidae) (blue dotted lines) after Luque et al. (2021).

For each vetted calibration, we reassess the earliest occurrences of several key groups in light of recent fossil findings and re-examination of previous ones. In some cases, the selection of a specific fossil calibration point from the literature does not necessarily represent the oldest record that has been putatively assigned to a given family, but the oldest reliable occurrence that can be referred to the clade with optimal confidence in its morphology. Minimum and tip maximum ages were either obtained from the ages reported in the original publications, or from stratigraphically constrained absolute or relative ages from the same localities as the fossil material or localities nearby, whenever possible. Numerical ages for geological time intervals follow the International Chronostratigraphic Chart by the International Commission on Stratigraphy (Cohen et al., 2013; updated 2023).

The identification and critical examination of key fossils occurrences is crucial to disentangle the anatomical disparity of crabs and their phylogenetic relationships, to recognize the lower limits of Brachyura, the origins of carcinization and decarcinization, the transition from sea to land and freshwater, and thus to ensure not only precise but more accurate time calibrations—both as tips and nodes—and divergence time estimates when reconstructing the crab tree of life.

## FOSSIL CALIBRATIONS

### 1. Anomura: Galatheoidea: Porcellanidae (crown)

#### Fossil specimen

*Vibrissalana jurassica* Robins and Klompmaker, 2019. Naturhistorisches Museum Wien, Vienna, Austria. Holotype NHMW 2007z0149/0405, a somewhat complete dorsal carapace, with a broken left side (Fig. 2A).

**Figure 2.**
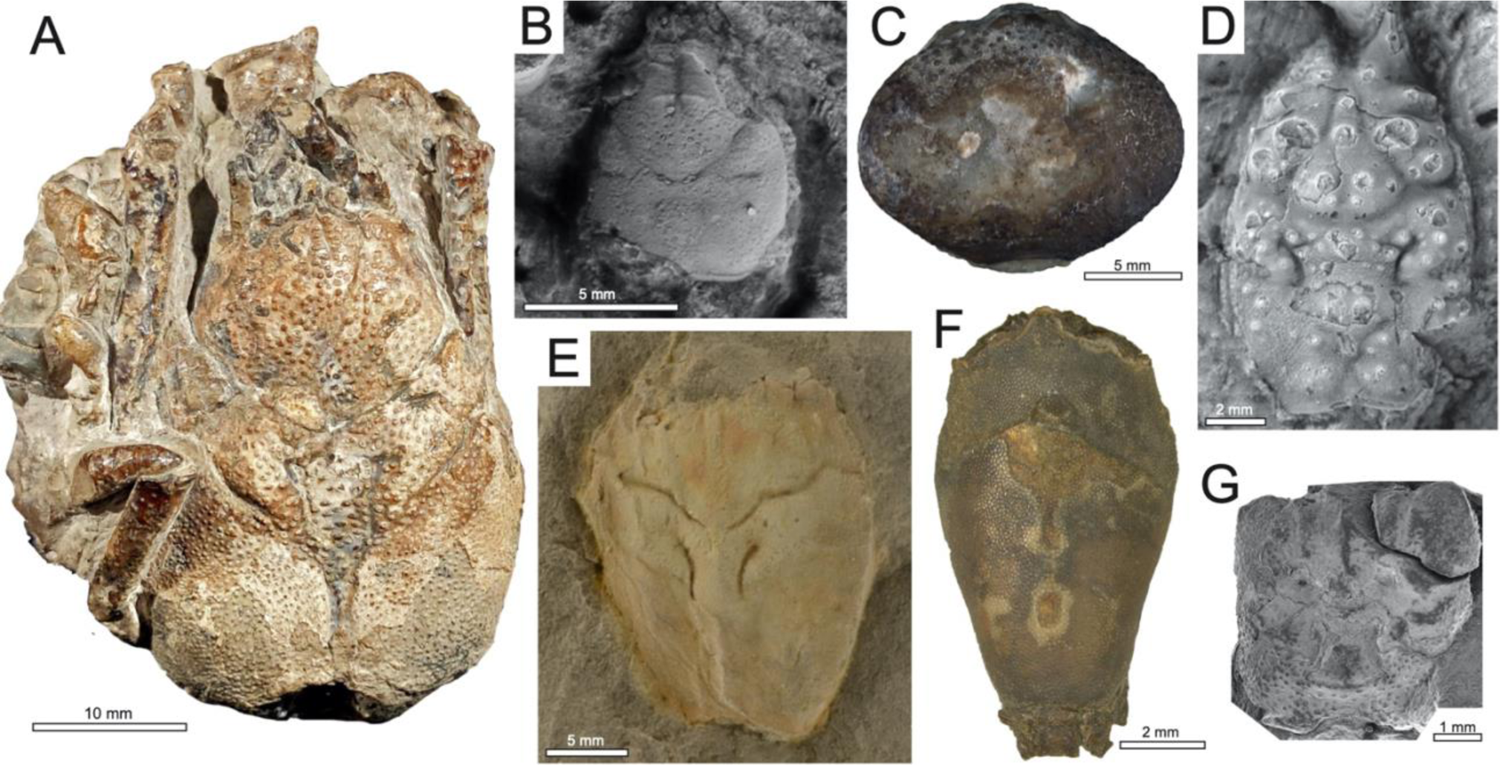
Meiura: **A**: *Eocarcinus praecursor* Withers, 1932, holotype NHM 18425, Lower Jurassic (Pliensbachian), UK. **B**: Anomura: **B**: *Vibrissalana jurassica* Robins and Klompmaker, 2019, holotype NHMW 2007z0149/0405, Upper Jurassic (Tithonian), Austria. **C–G**: Brachyura; **C**: Dromiacea: Dromioidea: *Graptocarcinus muiri* Stenzel, 1944, holotype BEG21288, Lower Cretaceous (Albian), San Luis Potosí, Mexico; **D**: Homoloida: Homoloidea: Homolidae: *Doerflesia ornata* Feldmann and Schweitzer, 2009, holotype Coll. No. 210, uppermost Lower Cretaceous (upper Albian), Texas, USA; **E–F**: Raninoida; **E**: Palaeocorystoidea: Palaeocorystidae: *Joeranina kerri* (Luque, Feldmann, Schweitzer, Jaramillo, and Cameron, 2012, as *Notopocorystes*), holotype IGM p881128, upper Lower Cretaceous (Aptian), Colombia; **F**: Raninoidea: Lyreididae: *Marylyreidus punctatus* (Rathbun, 1935, as *Notopocorystes*), non-type specimen Coll. No. 210 Texas, USA; **G**: Cyclodorippoida: Cyclodorippoidea: Cymonomidae: *Cymonomus primitivus* Müller and Collins, 1991., holotype EK-10.1 [M.91-135], Eocene (Priabonian), Budapest, Hungary. Photos by: Javier Luque (A, E, and F), Adiel Klompmaker (B), Ann Molineux, courtesy of Lisa Boucher and Adiel Klompmaker (C), Rodney Feldmann (D), and Alfréd Dulai (G).

#### Phylogenetic justification

The overall carapace shape of the holotype and sole specimen of *V. jurassica* fits well within the general diagnosis of Galatheoidea. The ovate, wide, and flattened dorsal carapace, plus the limited dorsal ornamentation and groove pattern, match previous diagnoses for Porcellanidae (Ahyong et al., 2010; Schweitzer and Feldmann, 2012; Robins and Klompmaker, 2019).

***Minimum age***. 145.0 Ma.

***Tip maximum age***. 152.0 Ma.

***Soft maximum age***. 191.8 Ma.

#### Age justification

This fossil was collected from the Ernstbrunn Limestone, exposed in the Ernstbrunn Quarries, Ernstbrunn, Austria. Although the age of the Ernstbrunn Limestone, based on microfossil and ammonite biostratigraphy, has been considered to range from the middle Tithonian (Upper Jurassic) to the Berriasian (Lower Cretaceous) (e.g., Moshammer and Schlagintweit, 1999; Zeiss, 2001; Schneider et al., 2013), the age of the rocks bearing meiurans has been consistently treated as middle upper Tithonian (*Richterella richteri* Zone to *Micracanthoceras microcanthum* Zone, *Simplisphinctes* Subzone) (e.g., Zeiss, 2001; Feldmann and Schweitzer, 2009; Schweitzer and Feldmann, 2010a; Robins et al., 2013; Fraaije et al., 2019; Robins and Klompmaker, 2019), for which a tip maximum age and a minimum age for *Vibrissalana jurassica* can be bracketed between ∼152.0 and 145.0 Ma.

As described in Wolfe et al. (2019), a soft maximum age is obtained by phylogenetic bracketing. The oldest crown Brachyura is debatable; *Eocarcinus praecursor* Withers, 1932, and *Eoprosopon klugi* Förster, 1986, have both been proposed, but both lack some crown group characters. Nevertheless, stem lineage positions of these taxa (Scholtz, 2020) allow a calibration of crown group Meiura with a soft maximum from the base of the Pliensbachian, at 191.8 Ma.

#### Discussion

*Jurellana tithonia* Schweitzer and Feldmann, 2010a, also from the Ernstbrunn Limestone, was previously considered the oldest porcellanid, although a recent revision of the taxon has suggested an affinity with homolodromioid brachyurans (Robins and Klompmaker, 2019). As such, *Vibrissalana jurassica* stands as the putatively oldest crown porcellanid (∼152–145 Mya) and represents, to date, a better calibration point for the crown group Porcellanidae.

### 2. Brachyura: Dromiacea: Dromioidea: Dynomenidae (crown)

#### Fossil specimen

*Graptocarcinus muiri* Stenzel, 1944. Bureau of Economic Geology, University of Texas at Austin. Holotype BEG00021288.000, a large and complete dorsal carapace (Fig. 2C).

#### Phylogenetic justification

The overall ovate carapace outline, being wider than long, with discrete but low-relief dorsal grooves, fits the diagnosis of Dynomenidae (Schweitzer et al., 2012; Van Bakel et al., 2012a).

***Minimum age***. 100.5 Ma.

*Tip maximum age.* 113.0 Ma.

*Soft maximum age.* 191.8 Ma.

#### Age justification

This fossil was collected in the Taniniúl limestone, upper Lower Cretaceous (Albian), from Choy Cave in Sierra del Abra between Las Palmas and Taninil, at kilometre 550 on the railroad between Tampico and San Luis Potosí, State of San Luis Potosí, Mexico (Stenzel, 1944).

Soft maximum age as for calibration 1.

#### Discussion

Six species of the genus *Graptocarcinus* Roemer, 1887, ranging in age from the Lower Cretaceous (Albian) to the middle Eocene (lower Lutetian) (Schweitzer et al., 2012; Beschin et al., 2016b), are currently included within the subfamily Graptocarcininae Van Bakel, Guinot, Corral & Artal, 2012a. Two isolated dactyli from the Barremian “Coulés boueuses”, Serre de Bleyton, France, have been adjudicated to *Graptocarcinus* (Hyzny and Kroh, 2015). Due the fragmentary nature of the samples, we considered for calibration the earliest occurrence of fossils with more diagnostic features that would warrant inclusion within crown Dynomenidae.

Several taxonomic studies have highlighted the issues of synonymy among some species within the genus *Graptocarcinus*, for instance, *Graptocarcinus muiri*, from the Albian of San Luis Potosí, Mexico, and *Graptocarcinus texanus* Roemer, 1887, from the Albian and Cenomanian of the US and Europe, with some of them concluding that *G. miuri* is a junior synonym of *G. texanus* (e.g., Klompmaker, 2013; Kocova-Veselská et al., 2014, and references therein), whereas other studies have maintained them as separate, valid species (e.g., Schweitzer et al., 2012; Van Bakel et al., 2012a; Beschin et al., 2016b, and references therein). Since the earliest confirmed occurrences of swell-preserved fossils of both species are Albian in age, choosing one species over the other would impact minimally the time calibration bracket for the genus (113–100 Mya).

## 3. Brachyura: Dromiacea: Dromioidea: Dromiidae (crown)

### Fossil specimen

*Dromiopsis gigas* Forir, 1887. Mineral collections of the Université de Liège, Belgium. Holotype 4936, a large, fragmented dorsal carapace in volume, right side.

### Phylogenetic justification

*Dromiopsis* is an extinct genus with over a dozen species known to date (Schweitzer et al., 2010). The overall circular carapace shape nearly as wide as long, the fronto-orbital configuration, and the well-developed set of dorsal grooves delimiting the dorsal carapace regions, guarantee placement within the crown Dromiidae (Schweitzer et al., 2012).

*Minimum age*. 93.9 Ma.

*Tip maximum age.* 100.5 Ma.

*Soft maximum age.* 191.8 Ma.

### Age justification

The type material of *Dromiopsis gigas* comes from the Tourtias (Formation?), Tournai, Belgium, which has been dated as Cenomanian based on ammonite biozonation (Kennedy et al., 2011).

### Discussion

*Costadromia hajzeri* Feldmann and Schweitzer, 2019, from the Campanian (Upper Cretaceous, ∼83–72 Mya) Wenonah Formation in New Jersey, USA, has been recently proposed as the earliest known sponge crab, and included within Dromiidae sensu lato based mostly on details of the frontal margin (Feldmann and Schweitzer, 2019). While *C. hajzeri* does seem to fit well within the total group Dromiidae, the presence of *Dromiopsis gigas* in mid-Cretaceous (Cenomanian, ∼100–93 Mya) rocks of Belgium, and its conspicuous similarity to other genera within crown Dromiidae, indicate that the type material of *D. gigas* provides a better current calibration point for crown Dromiidae.

Soft maximum age as for calibration 1.

## 4. Brachyura: Homoloida: Homoloidea: Homolidae (stem)

### Fossil specimen

*Doerflesia ornata* Feldmann and Schweitzer, 2009. Naturhistorisches Museum Wien (NHMW). Holotype 2007z0149/0015, a small dorsal carapace nearly complete (Fig. 2D).

### Phylogenetic justification

The genus *Doerflesia* has several of the diagnostic features of the family Homolidae, and it shows some resemblance to extant genera (Feldmann and Schweitzer, 2009; Feldmann et al., 2012). While a homolid affinity is most likely, its position within the crown group cannot be warranted with the available fossil material (a dorsal carapace of the holotype and sole specimen), therefore we have chosen to treat it as a stem group member of Homolidae.

***Minimum age***. 145.0 Ma.

*Tip maximum age.* 152.0 Ma.

*Soft maximum age.* 191.8 Ma.

### Age justification

This fossil was collected from the Ernstbrunn Limestone, exposed in the Ernstbrunn Quarries, near the village of Dörfles, Austria.

Soft maximum age as for calibration 1.

## 5. Brachyura: Raninoida: Palaeocorystoidea: Palaeocorystidae (stem)

### Fossil specimen

*Joeranina kerri* (Luque, Feldmann, Schweitzer, Jaramillo, and Cameron, 2012) (as ss*Notopocorystes*). Colombian Geological Survey (formerly INGEOMINAS), Museo Geológico José Royo y Gómez, Bogotá DC, Colombia. Holotype IGM p881128, a nearly complete dorsal carapace (Luque et al., 2012, pp. 411–413, fig. 4A, B.; Luque et al., 2017, p. 20, fig. 8B) (Fig. 2E).

### Phylogenetic justification

Recent phylogenetic studies have shown consistently that crabs of the extinct superfamily Palaeocorystoidea occupy an intermediate place between the crab-like Necrocarcinoidea and the frog-like Raninoidea (Karasawa et al., 2014; Luque, 2015a; Schweitzer et al., 2016a; Luque et al., 2019b). As such, Palaeocorystoidea as a whole is recovered as part of the stem group of Raninoidea, the latter of which includes all of the extant genera of frog crabs.

***Minimum age***. 113.0 Ma.

*Tip maximum age.* 118.0 Ma.

*Soft maximum age.* 191.8 Ma.

### Age justification

The holotype of *Joeranina kerri* was discovered *in-situ* by one of us (J.L.) in grey shales of the upper portion of the Lower Cretaceous Paja Formation, exposed along the road between the towns of San Gil and Curití, in the Department of Santander, Colombia. The occurrence of the gastropod *Turritella* (*Haustator*) *columbiana* Jaworski, 1938, and the ammonite *Acanthohoplites eleganteante* Etayo-Serna, 1979, stratigraphically below the horizon yielding the holotype of *J. kerri*, indicate an upper Aptian age in Colombia (see in Luque et al., 2012; Luque, 2014).

Soft maximum age as for calibration 1.

### Discussion

Frog crabs and allies, together constituting the Section Raninoida (Ahyong et al., 2007), are among the most anatomically disparate brachyuran groups (Schweitzer et al., 2012; Van Bakel et al., 2012b; Karasawa et al., 2014; Hartzell et al., 2022). Recent phylogenetic studies have shown that early-branching forms, referred to as the crab-like raninoidans (e.g., Orithopsidae, Paranecrocarcinidae, Necrocarcinidae, Cenomanocarcinidae) are sister groups to a clade formed by mostly decarcinized, frog-like raninoidan families such as the extinct Palaeocorystidae and the extant Raninidae and allied relatives (Luque, 2015a; Schweitzer et al., 2016a; Luque et al., 2019b).

As such, the most recent common ancestor of Palaeocorystidae + Raninidae must be as old or older than the oldest fossils known within Palaeocorystidae, which correspond to *Joeranina kerri* from the upper Aptian (Lower Cretaceous, 118–113 Mya) of Colombia, South America, as indicated above.

## 6. Brachyura: Raninoida: Raninoidea: Lyreididae (stem)

### Fossil specimen

*Marylyreidus punctatus* (Rathbun, 1935, as *Notopocorystes*). University of Texas. Holotype Coll. No. 210, a nearly complete dorsal carapace in volume. A non-type specimen is illustrated in Fig. 2E.

### Phylogenetic justification

The type material of *Marylyreidus punctatus*, together with additional specimens known from different localities and preserving dorsal, ventral, cuticular, and pleonal details, strongly show a crown Raninoidea affinity, with a sternal configuration suggesting a close proximity to lyreidids (Van Bakel et al., 2012b; Karasawa et al., 2014; Frantescu et al., 2016; Schweitzer et al., 2018). This is consistent with the phylogenetic position of *Marylyreidus*, recovered in a parsimony analysis as sister to *Bournelyreidus* van Bakel, Guinot, Artal, Fraaije, and Jagt, 2012b, both forming an early diverging stem group sister to the remainder of the lyreidid-like frog crabs (Karasawa et al., 2014).

***Minimum age***. 100.5 Ma.

*Tip maximum age.* 104.0 Ma.

*Soft maximum age.* 191.8 Ma.

### Age justification

The type specimen of *Marylyreidus punctatus* was discovered in rocks of the Lower Cretaceous (Albian), indicated by Rathbun (1935) as belonging to the ‘Denton Clay, lower Comanche Series, Washita Formation’, cropping out in Grayson County, two miles north of Denison, Texas, USA (Rathbun, 1935). Currently, the Washita ‘Formation’ in Texas is considered at the Group level, and constituted at its thickest part by seven formations, in ascending order: Kiamichi, Duck Creek, Fort Worth, Denton, Weno, Paw Paw, and Main Street formations (Scott et al., 2002; Scott et al., 2016). The entire Washita Group from base to top is thought to have been deposited between 104–94.4 Ma, with its lower boundary in the lowermost upper Albian, and with the upper Albian–lower Cenomanian boundary (100.5 Ma) represented in the uppermost Main Street Limestone (Scott et al., 2000; Scott et al., 2002). Denton Formation has been dated as upper Albian, based on the occurrence of ammonites from the *Mortoniceras* (*Subschloenbachia*) *rostratum* Zone (Kennedy et al., 2005). Additional specimens assigned to *M. punctatus* have been recovered from several localities and stratigraphic intervals in Texas, including the slightly younger but also upper Albian Paw Paw Formation (Frantescu et al., 2016). As such, relaxing the lowermost and uppermost potential occurrences of *M. punctatus* within the Washita Group, its tip maximum and minimum ages can be bracketed within 104.0–100.5 Ma.

Soft maximum age as for calibration 1.

### Discussion

Wolfe et al. (2022) recovered a monophyletic Lyreididae nested within a paraphyletic ‘Raninidae’. As such, despite *Marylyreidus* being part of the stem group to Lyreididae (or Lyreininae, if confirmed to be a subfamily within Raninidae), it belongs to the crown Raninoidea. Besides *M. punctatus*, only a couple other raninoideans are currently known from upper Albian rocks worldwide, both initially assigned to the genus *Hemioon* Bell, 1863, traditionally considered a Lyreididae. These are ‘*H. cunningtoni’* Bell, 1863, the type species of the genus and known from the upper Albian of England, and ‘*Hemioon yanini*’ Ilyin and Alekseev, 1998, known from the upper Albian of Crimea. However, both records are problematic. Firstly, ‘*Hemioon*’ has been a problematic taxon due to the few specimens known and their poor preservation, resulting in some authors considering it a valid taxon, whereas others have synonymized it with *Raninella* A. Milne-Edwards, 1862a (see discussions in Bishop and Williams, 2000; Van Bakel et al., 2012b; Karasawa et al., 2014; Schweitzer et al., 2018 and references therein). Secondly, *Raninella* ‘*cunningtoni*’ might be a juvenile of *Raninella elongata* Milne-Edwards, 1862b (see Bishop and Williams, 2000; Van Bakel et al., 2012b; Karasawa et al., 2014 and references therein). Thirdly, *Raninella* is currently considered a member of the subfamily Ranininae within the Raninidae, in which case this family would also have an upper Albian record. Unfortunately, besides the conflicting views on their systematic affinities, the exact age of ‘*H. cunningtoni*’ needs further revision (B. van Bakel, pers. comm. to J.L., February 2023).

Regarding ‘*Hemioon yanini*’, its age is better constrained the species above discussed, being referable to the upper Albian based on its occurrence in the *Mortoniceras infiatum* zone (Ilyin and Alekseev, 1998; Ilyin, 2005), but its systematic placement is equally convoluted. Van Bakel et al. (2012b) included this species within *Raninella*, whereas Karasawa et al. (2014) included it within the genus Macroacaena Tucker, 1998, under the lyreidid subfamily Macroacaeninae Karasawa, Schweitzer, Feldmann, and Luque, 2014 (Schweitzer et al., 2018). Regardless the generic and specific affinities of ‘*H. cunningtoni*’ and ‘*H. yanini*’, the presence of disparate raninidoid forms in the late Albian of USA, UK, and Crimea, indicate that the most recent common ancestor of Raninodiea and all of its descendants must have originated before the late Albian.

## 7. Brachyura: Cyclodorippoida: Cyclodorippoidea: Cymonomidae (crown)

### Fossil specimen

*Cymonomus primitivus* Müller and Collins, 1991. Természcttudományi Múzeum, Föld-és oslénytar, VIII Muzeum krt. 14-16, H-1088 Budapest, Hungary. Holotype EK-10.1 [M.91-135], a partial dorsal carapace (Müller and Collins, 1991, p. 61, 63–64, pl. 3, fig. 6) (Fig. 2G).

### Phylogenetic justification

The Hungarian specimen can be referred to crown Cyclodorippoidea, and assignable to Cymonomidae based on diagnostic characteristics of the subquadrate dorsal carapace, the narrow fronto-orbital margin, and the configuration of the dorsal grooves and regions, which fit well within the diagnosis of crown Cymonomidae (Müller and Collins, 1991; Tavares, 1993; Ahyong, 2019).

*Minimum age*. 33.9 Ma.

*Tip maximum age.* 37.71 Ma.

*Soft maximum age.* 191.8 Ma.

### Age justification

*Cymonomus primitivus* was collected from 4–5 m thick coral-bearing limestones of Facies 4 of the Upper Eocene (Priabonian) Szépvölgy Limestone Formation, cropping out at the Ruprecht quarry, Budapest, Hungary, with the coral fauna indicating a Priabonian age (Müller and Collins, 1991).

Soft maximum age as for calibration 1.

### Discussion

Besides *Cymonomus primitivus*, other presumed cymonomids are known from the Eocene, e.g., *Spathanomus felicianensis* De Angeli, 2016, and *Caporiondolus bericus* De Angeli, 2016, from the Priabonian of of the Orgiano quarry, Monti Berici, Italy, and *Eonomus californianus* Nyborg, Garassino, and Slak, 2017, from the early to middle Eocene LLajas Formation, Simi Valley, California, USA. We selected *C. primitivus* as our vetted calibration point for crown Cymonomidae (∼37–33 Ma), since *Cymonomus* is an extant genus, whereas *Spathanomus*, *Caporiondolus*, and *Eonomus* are all extinct genera, and their family-level affinities are considered by some authors as uncertain (e.g., Schweitzer et al., 2017).

## 8. Brachyura: Eubrachyura: Potamoidea: Potamidae (crown)

### Fossil specimen

*Alontecarcinus buratoi* De Angeli and Caporiondo, 2019. Museo di Storia Naturale di Verona. Holotype IGVR 19.38, a complete dorsal carapace (Fig. 3B).

**Figure 3.**
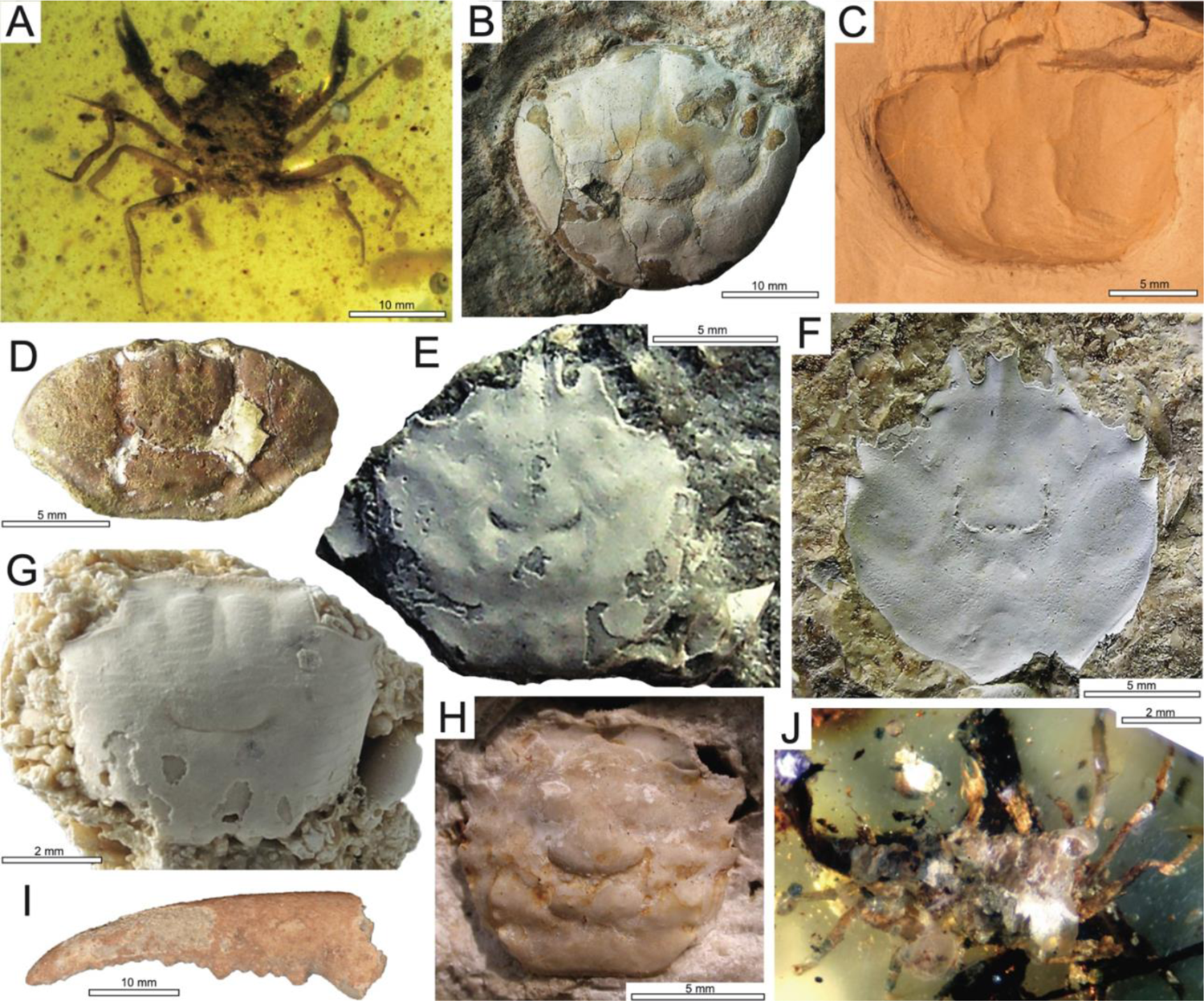
Crown Eubrachyura. **A**: Superfamily uncertain: Cretapsaridae: *Cretapsara athanata* Luque in Luque et al., 2021, holotype LYAM-9, lowermost Upper Cretaceous (lower Cenomanian), Burma. **B**: Potamoidea: Potamidae: *Alontecarcinus buratoi* De Angeli and Caporiondo, 2019, holotype IGVR 19.38, Eocene (Bartonian), Vicenza, Italy. **C– J**: Eubrachyura: Thoracotremana; **C**: Ocypodoidea: Ocypodidae: *Afruca miocenica* (Artal, 2008, as *Uca*), holotype MGSB 68653, Miocene (Langhian), Catalonia, NE Iberian Peninsula. **D**: Pinnotheroidea: Pinnotheridae: *Pinnixa* sp., specimen UF 115397, lower Miocene (Burdigalian), Panama Canal, Panama (Luque et al., 2017). **E–J**: Grapsoidea; **E,F**: Percnidae; **E**: *Percnon santurbanensis* Ceccon and De Angeli, 2019,, holotype MCV.19/02, lower Oligocene (Rupelian), Vicenza, Italy; **F**: *Percnon paleogenicus* De Angeli, 2023., holotype MCV.23/738-22.341, upper Eocene (Priabonian) Vicenza, Italy. **G**: Grapsidae: *Pachygrapsus hungaricus* Müller, 1974, holotype HNHM 2004.163.1, middle Miocene (Badenian), Budapest area, Hungary. **H**: Varunidae: *Brachynotus corallinus* Beschin, Busulini, De Angeli & Tessier, 2007, holotype MCZ 1794, lower Eocene (Ypresian), Vicenza, Italy. **I**: Gecarcinidae: *Cardisoma guanhumi* Latreille, in Latreille, Le Peletier, Serville and Guérin, 1828, specimen XXXX, Pleistocene, XXXXX. **J**: Sesarmidae indet., specimen IHNFG-4991, lower Miocene (Aquitanian) of Chiapas, Mexico (Serrano-Sánchez et al., 2016). Photos by: Lida Xing (A), Antonio de Angeli (B, E, F), Pedro Artal (C), Javier Luque (D, I), Matus Hyzny (G), Alessandra Busulini (H), and Francisco Vega (J).

### Phylogenetic justification

The type material of *Alontecarcinus buratoi* shows remarkable similarities with the dorsal carapaces of extant potamid crabs, including their oval outline and wide carapace, the well-developed cervical and gastro-cardiac grooves, a front that is entire and directed downwards, and the convex lateral margins that are smooth and bear a single epibranchial tooth (De Angeli and Caporiondo, 2019).

***Minimum age***. 37.71 Ma.

*Tip maximum age.* 41.2 Ma.

*Soft maximum age.* 100.9 Ma.

### Age justification

This fossil was collected from middle Eocene limestones of the Alonte quarry in Berici Mounts, Vicenza, Northern Italy (De Angeli and Caporiondo, 2019). The stratigraphic position of the Alonte quarry Limestones, overlying rocks of middle Eocene age, and underlying rocks of Priabonian age, together with associations of gastropods and bivalves (e.g., *Ampullina*, *Cerithium*, *Campanile*, *Natica*, *Corbis*, *Glycimeris*), echinoids (e.g., *Leiopedina*, *Sismondia*, *Echinolampas*, *Schizaster*, *Cidaris*), and calcareous nannofossils, indicate a Bartonian age for the Alonte quarry Limestones (41.2–37.71 Ma) (Beccaro, 2003; De Angeli and Alberti, 2016; De Angeli and Caporiondo, 2019).

As this node is within Eubrachyura, a soft maximum age is based on *Cretapsara athanata* Luque in Luque et al., 2021, which is the oldest crown group Eubrachyura. The position of *C. athanata* relative to modern eubrachyuran families is not clear, but its membership within the crown group, based on morphological phylogenetic analysis (Luque et al., 2021), indicates that the common ancestor of typical eubrachyuran forms must be older than the earliest Late Cretaceous. The fossil was found in Burmese amber, for which the exact age estimates can vary, depending of the source. The limited radioisotopic U-Pb information available for Kachin burmite indicates an age of 98.8 ± 0.6 Ma (Shi et al., 2012), which would suggest a bracketed tip maximum age and minimum age for *C. athanata* between 99.4 Ma and 98.2 Ma. As radiometric estimates pertain primarily to the sediments and not necessarily the amber itself (see discussion in Luque et al., 2021). Therefore, we calibrate a soft maximum close to the Albian–Cenomanian boundary 100.5 Ma ± 0.4 Myr = 100.9 Ma.

## 9. Brachyura: Eubrachyura: Ocypodoidea: Ocypodidae (crown)

### Fossil specimens

*Afruca miocenica* (Artal, 2008, as *Uca*). Museo Geológico del Seminario de Barcelona (MGSB), Catalonia, Spain. Holotype MGSB 68653, a complete and well-preserved dorsal carapace (Fig. 3C), and paratypes 68654a to 68654e, including well-preserved dorsal and ventral males and females with pleon, maxillipeds, pereopods, and the male major cheliped.

### Phylogenetic justification

The type material of *Afruca miocenica* can be confidently assigned to the genus *Afruca* based on the diagnostic enlarged and flattened dactyli and pollices of the male major cheliped, and it is closest to the extant *Afruca tangeri* (Eydoux, 1835), which is the only recognized species of fiddler crabs living today in the Iberic Peninsula and north Atlantic Africa. While *A. miocenica* might indeed represent its own species, its overall similarities with *A. tangeri* have invited the question of whether they are synonymous, with the former representing ontogenetic variations similar to those seen in younger individuals of the later (de Gibert et al., 2013). In either scenario, the close proximity between *A. miocenica* and *A. tangerii* is remarkable, and the phylogenetic assignment of the fossils from Catalonia to the extant genus *Afruca*—and thus to the crown Ocypodidae—are confirmed.

***Minimum age***. 13.82 Ma.

*Tip maximum age.* 15.97 Ma.

*Soft maximum age.* 100.9 Ma.

### Age justification

The fossil material of *Afruca miocenica* was collected *in-situ* in terrigenous yellow claystones from a lower Middle Miocene (Langhian, 15.97–13.82 Ma) locality of the Rubí (Vallés), Province of Barcelona, Catalonia, NE Iberian Peninsula (Artal, 2008). These deposits are part of the Vallès-Penedès basin, and represent marine to coastal mangrove facies deposited during the maximum transgression of the Langhian in the area (Batllori and García, 1997; de Gibert and Robles, 2005; Artal, 2008; Garassino et al., 2009), overlying continental Burdigalian facies (Casanovas-Vilar et al., 2011), and underlying continental Serravalian facies (Batllori and García, 1997; de Gibert and Robles, 2005; Casanovas-Vilar et al., 2016).

Soft maximum age as for calibration 8.

### Discussion

The fossil record of fiddler crabs is sparse and fragmentary, with only half a dozen species recognized from fossilized carapaces (e.g., Artal, 2008; Domínguez Alonso, 2008; de Gibert et al., 2013; Luque et al., 2017; Luque et al., 2018; Lima et al., 2020). Aside *Afruca miocenina*, the only other Miocene fiddler crab known is *Uca maracoani* (Latreille, 1802) from Brazil (Brito, 1972; Luque et al., 2017; Luque et al., 2018; Lima et al., 2020). The fossil material of *U. maracoani* was collected in rocks of the Pirabas Formation, in the village of Baunilha Grande, Baia de Quatipuru, Pará State, Brazil (Brito, 1972), and subsequent material from the area has been reported recently (Lima et al., 2020). However, the exact age and stratigraphic provenance of these specimens is unclear. Overall, palynological, lithological, petrographic, and geochemical data indicate that the Pirabas Formation as a whole is constituted by two distinct depositional groups: the oldest and lowermost one, of uppermost Lower Miocene age, deposited in a shallow marine environment, and the youngest and uppermost one, of upper Middle Miocene to Upper Miocene, deposited in a coastal environment near mangroves (e.g., Aguilera et al., 2022; Gomes et al., 2023). In localities such as Capanema, where the lowermost depositional unit outcrops, absolute Sr/Sr dating obtained from pectinid bivalve shells have yielded ages of 17.3–16.0 Ma, suggesting an uppermost Lower Miocene (upper Burdigalian) age (Martinez et al., 2017; Gomes et al., 2023). In localities such as Praia do Atalaia and Praia do Maçarico, where the uppermost depositional unit outcrops, palynomorphs of the T16 biozone (Jaramillo et al., 2011) indicate an age of 12.7 to 7.1 Ma, suggesting an uppermost Middle Miocene to Upper Miocene age (mid Serravalian to lowermost lower Messinian) (Gomes et al., 2023). The fiddler crab-bearing facies of the Pirabas Formation at the ‘Furo de Baunilha Grande’ locality correspond to the latter, younger unit, with a most probable age of uppermost Middle Miocene to Upper Miocene, roughly corresponding to the Serravalian–Tortonian (13.82–7.246 Ma) (Gomes et al., 2023). Moreover, these fiddler crab-bearing nodules are not *in-situ*, but found loose on the Baunilla stream floor (Lima et al., 2020; Orangel Aguilera, pers. comm. to J.L., February 2023), and although previous works have presumably recovered palynological samples associated to these nodules that would suggest a uppermost Lower Miocene rock age (e.g., Antonioli et al., 2015), more recent and comprehensive biostratigraphic and paleonvironmental studies across localities have confirmed the uppermost Middle Miocene to Upper Miocene age of the Baunilha Grande rocks (Gomes et al., 2023). As such, we opt to use the apparently older and *in-situ* occurrence of *Afruca miocenica* (Langhian, 15.97–13.82 Ma) as our calibration point for Ocypodidae.

Aside from the two fiddler crab species above mentioned, the only other Ocypodidae with Miocene fossils known to date is *Ocypode vericoncava* Casadío et al., 2005, a single damaged dorsal carapace from the upper Miocene (Tortonian? 11.63–7.246 Ma) of Argentina. Despite the poor preservation of the holotype and sole specimen, the overall shape and fronto-orbital configuration match those of *Ocypode*. Thus, this occurrence from Argentina, together with those of *Uca* from Brazil and *Afruca* in Catalonia, indicate that Ocypodidae was already widespread by the mid-Miocene, and that the most recent common ancestor of Ocypodidae and its divergence from other thoracotremes must have occurred in the pre-Neogene, most likely in the Paleogene or even the Cretaceous (Wolfe et al., 2022).

## 10. Brachyura: Eubrachyura: Pinnotheroidea: Pinnotheridae (crown)

### Fossil specimen

*Pinnixa* sp. Invertebrate Paleontology collections, Florida Museum of Natural History. Specimen UF 115397, a complete dorsal carapace (Luque et al., 2017, fig. 12Q) (Fig. 3D).

### Phylogenetic justification

Based on the overall carapace shape and dorsal patterns, this fossil specimen can be assigned with confidence to the genus *Pinnixa*. Recent molecular studies, however, have shown that *Pinnixa* species may not form a monophyletic genus, but be distributed among three Pinnotheridae subfamilies (e.g., Palacios Theil et al., 2016). Despite this, they all can be assigned to the crown-group Pinnotheridae, justifying the inclusion of the fossil *Pinnixa* from the lower Miocene of Panama in the crown group.

*Minimum age*. 18.7 Ma.

*Tip maximum age.* 21.68 Ma.

*Soft maximum age.* 100.9 Ma.

### Age justification

The specimen was collected *in-situ* by one of us (J.L.) in marine grey mudstones from the middle part of the upper member of the Culebra Formation in the Panama Canal Zone, Lirio section, Culebra Cut (formerly known as Gaillard Cut). The marine rocks of the Culebra Formation are lower Miocene (uppermost Aquitanian–lower Burdigalian), and not Oligocene as assumed by Rathbun, 1918 [1919] and repeated as such by subsequent authors (see notes in Luque et al., 2017). At its base, the marine Culebra Formation overlies in sharp unconformity the terrestrial upper Oligocene-lower Miocene rocks of the Las Cascadas Formation, and its top, it underlies conformably the coarser terrestrial rocks of the upper Lower to Middle Miocene Cucaracha Formation (Montes et al., 2012; Buchs et al., 2019; LeBlanc, 2021). Kirby et al. (2008) reported Sr isotope dating ages for different parts of the formation, including four ages from *Acropora* corals and pectinids in the middle member of the Culebra Formation (Emperador Limestone), ranging between 21.24±0.44 and 20.62±0.58 Ma, and two ages of from a pectinid and an ostreid (19.83±0.39 and 19.12±0.42 Ma, respectively), from the lowermost portion of the Cucaracha Formation, above the top of the upper member of the Culebra Formation (Kirby et al., 2008, fig. 6, table 2). Montes et al. (2012) reported a U/Pb age of 19.3 ±0.4 Ma from magmatic zircons in an ash tuff from the lower half of the upper member of the Culebra Formation, and MacFadden et al. (2014) reported U/Pb and Ar/Ar ages of 18.81±0.3 and 18.96±0.9, respectively, from detrital zircons in an ash tuff from the overlying Cucaracha Formation. As such, based on the currently available absolute ages available, the age of the crab-bearing beds in the middle part of the upper member of the Culebra Formation— and thus that of *Pinnixa* sp.—can be bracketed with confidence between 21.68 and 18.7 Ma, most likely around 20–19 Ma (see Farris et al., 2017), in the lowermost Burdigalian, near the Aquitanian–Burdigalian boundary.

Soft maximum age as for calibration 8.

### Discussion

The fossil record of pinnotherids is among the most controversial among eubrachyurans, in part because of their overall small sizes, their largely fragmentary fossil record, and the overall carapace outline similarities of some fossils to some species of other eubrachyuran (e.g., some hexapodids and aphanodactylids). This obscure the systematic affinities of several fossils, and thus cast doubts of their usefulness for node calibrations of crown Pinnotheridae.

The extinct genus *Viapinnixa* Schweitzer and Feldmann, 2001, comprises four species that range in age from the lower Paleocene to the lower Eocene (e.g., Philippi, 1887; Vega et al., 2001; Vega et al., 2007; Vega et al., 2008; Armstrong et al., 2009), which are remarkably similar to species within Hexapodidae. As such, the genus cannot be assigned with certainty to the pinnotheroids (Luque et al., 2017), and even less so to the crown group Pinnotheridae. Other fossil currently placed in the family Pinnotherinae include the spectacularly preserved extinct genus *Pharkidodes* Feldmann, Schweitzer, Casadío & Griffin, Feldmann et al., 2011b, from the middle Miocene of Tierra del Fuego, Argentina, which unfortunately cannot be placed within the crown group with certainty.

Three fossil species have been dubiously assigned to *Pinnotheres*, i.e., *P.*? *araucana* Philippi, 1887, from the ‘Tertiary’ of Chile, *P.*? *elatus* Milne-Edwards, 1873, from the upper Miocene of France, and *P.*? *promaucanus* Philippi, 1887, from the Miocene of Chile (Schweitzer et al., 2010). Over half a dozen species have been previously assigned to *Pinnixa*. ‘*Pinnixa*’ *eocenica* Rathbun, 1926, from the Eocene of Washington, USA, is a junior homonym of *Pinnixa eocenia* Woods, 1922, and it is currently recognized as a hexapodid: *Palaeopinnixa rathbunae* Schweitzer, Feldmann, Tucker & Berglund, 2000.

*Pinnixa navidadensis* Feldmann, Schweitzer and Encinas, 2005, from the middle Miocene of Chile (Feldmann et al., 2010; Jagt et al., 2015; and *Pinnixa* sp., from the middle Pleistocene of Florida (Agnew, 2001; Portell and Agnew, 2004), can be assigned to *Pinnixa*, but they are younger than *Pinnixa* sp. from the lower Miocene Culebra Formation in Panama. *Pinnixa aequipunctata* Morris and Collins, 1991, and *P. omega* Morris and Collins, 1991, come from the Pliocene Upper Miri Formation, Brunei. *Pinnixa microgranulosa* Collins et al., 2003, from the Miocene Sandakan Formation, Sarawak, may also belong to the hexapodid genus *Palaeopinnixa*.

As such, given the reliable systematic placement and the constrained stratigraphic age of *Pinnixa* sp. from the lower Miocene Culebra Formation in the Panama Canal Zone (Luque et al., 2017), this is to date the most reliable calibration datum for the crown group Pinnotheridae, while the most recent common ancestor of all extant pinnotherids must have originated in the pre-Neogene, most likely the Paleogene or Late Cretaceous (see Wolfe et al., 2022).

## 11. Brachyura: Eubrachyura: Grapsoidea: Percnidae (crown)

### Fossil specimen

*Percnon paleogenicus* De Angeli, 2023. Museo Civico “D. Dal Lago” di Valdagno (MCV), Vicenza, northern Italy. Holotype MCV.23/738-22.341, a complete dorsal carapace, part and counterpart (Fig. 3E).

### Phylogenetic justification

Seven extant species in the genus *Percnon* constitute the monophyletic family Percnidae Stevcic, 2005, which is nested within Grapsoidea (e.g., Schubart and Cuesta, 2010; Wolfe et al., 2022). The fossil species *P. paleogenicus* has a carapace shape and an orbitofrontal configuration that are typical of Percnidae close to *Percnon*, and thus is a reliable calibration point for the family.

*Minimum age*. 33.9 Ma.

*Tip maximum age.* 37.71 Ma.

*Soft maximum age.* 100.9 Ma.

### Age justification

The holotype of *P. paleogenicus* was collected from calcarenites and calcareous algae levels of the Collina di San Feliciano (Orgiano, Vicenza), which has been dated as upper Eocene (Priabonian, ∼37.71–33.9 Mya) based on nannofossils (Beccaro, 2003, in De Angeli and Garassino, 2021b, a) (Fig. 1).

### Discussion

Until recently, the only and thus oldest fossil assignable to Percnidae was *Percnon santurbanensis* Ceccon and De Angeli, 2019, represented by a well-preserved dorsal carapace collected from the lower Oligocene (Rupelian, ∼33.90–27.82 Ma) of the Berici eastern Lessini mountains in Montecchio Maggiore, Vicenza, northern Italy (Ceccon and De Angeli, 2019) (Fig. 3E). We used this fossil recently in Wolfe et al., 2022 as the calibration point for Percnidae, since *P. paleogenicus* was published posterior to the completion of the phylogenetic study and became available to us at a later stage. What is clear is that there is more than one fossil species referrable to Percnidae and *Percnon* per se, and that the crown group Percnidae must have a pre-Priabonian origin, probably into the Paleocene or uppermost Cretaceous.

## 12. Brachyura: Eubrachyura: Grapsoidea: Grapsidae (crown)

### Fossil specimen

*Pachygrapsus hungaricus* Müller, 1974. Hungarian Natural History Museum, department of Earth Sciences and Palaeontology. Holotype HNHM 2004.163.1, a nearly complete dorsal carapace (Fig. 3G).

### Phylogenetic justification

Morphological and molecular phylomitogenomic studies indicate that *Pachygrapsus* is phylogenetically well nested within crown Grapsidae, close to *Grapsus* (e.g., Karasawa and Kato, 2001; Chen et al., 2019; Lü et al., 2022; Tsang et al., 2022; Wolfe et al., 2022; Zhang et al., 2022).

*Minimum age*. 12.6 Ma.

*Tip maximum age.* 13.82 Ma.

*Soft maximum age.* 100.9 Ma.

### Age justification

The type material of *Pachygrapsus hungaricus* was collected from a patch reef with *Porites* of the Leitha Limestone Formation, exposed at Téteny-plateau, western part of the Budapest area, Hungary, dated as Middle Miocene (late Badenian, Decapod Zone 4 sensu Müller, 1984) (Hyzny, 2016; Hyzny and Dulai, 2021). Several tens of additional specimens referred to *P. hungaricus* have also been reported from coeval rocks in several localities in the Budapest area (Gyakorló út, Örs vezér tere, and Rákos), as well as in its vicinity (i.e., Biatorbágy and Diósd) in Hungary, and from coeval strata in Austria and Poland (Müller, 1984, 1996; Hyzny, 2016; Hyzny and Dulai, 2021). Other conspecific occurrences are known from slightly older strata as well (Decapod Zone 3 sensu Müller, 1984). Thus*, P. hungaricus* has been reliably documented from the entire span of the upper Badenian (13.82–12.6 Ma) (Kovác et al., 2018; M. Hyzny, pers. comm. to J.L., February 2023) (Figs 1, 3G).

The age of the lower and upper boundaries of the Paratethyan Badenian stage/age have been a matter of ongoing discussion. While some authors have considered that the age of the Badenian lower boundary is positioned at the top of chron C5Cn.2n (16.303 Ma), roughly equivalent to the uppermost part of the global Lower Miocene Burdigalian stage/age (e.g., Piller et al., 2007; Hohenegger et al., 2014; Wagreich et al., 2014), others have considered it to be closer to the base of the global Langhian stage/age (15.97 Ma) (Krijgsman and Piller, 2012; Reichenbacher et al., 2013; Gozhyk et al., 2015; Kovác et al., 2018). Similarly, for several authors, the age of the Badenian upper boundary is positioned at the top of polarity chron C5Ar.2n (12.829), roughly equivalent to the middle part of the global Middle Miocene Serravallian stage/age boundary (e.g.Hohenegger et al., 2014; Wagreich et al., 2014), whereas for others, the Badenian/Sarmatian boundary is slightly younger (12.7–12.6 Ma) (e.g., Piller et al., 2007; Krijgsman and Piller, 2012; Kovác et al., 2018, see also Hyzny and Dulai, 2021 and references therein). This reflects the issues of interpreting minimum and tip maximum ages for taxa based on the literature alone, since the lithostratigraphic boundary ages in different localities might vary in relation with eustatic sea-level changes and local subsidence, and rarely a crab fossil occurrence is tied to absolute dating.

Since the goal of our calibration is not to resolve the exact stratigraphic age of a given occurrence but to provide a confidence range that brackets the minimum and tip maximum ages, we opt to consider the minimum age of *P. hungaricus* as 12.6 Ma, and the tip maximum age as 13.82 Ma, which roughly delimit the bottom and top of the Late Badenian (Kovác et al., 2018).

Soft maximum age as for calibration 8.

### Discussionss

Fossils of grapsid crabs are rare, in part due to the relatively dynamic and high-energy environments they inhabit. Among grapsids, the genera *Goniopsis* and *Leptograpsus* (monotypic) have no confirmed fossil record. The genera *Geograpsus* and *Grapsus* have fossils known from the Holocene of Hawaii, i.e., *Geograpsus severnsi* Paulay and Starmer, 2011, and the Holocene of the Pacific in Panama, i.e., *G.* aff. *G. grapsus* (Luque et al., 2018). The extant genus *Metopograpsus* has two extinct species currently assigned to it, e.g., *M. badenis* Müller, 2006, from the same middle Miocene (Badenian) locality in Hungary as *Pachygrapsus hungaricus*, and *M. traxleri* Müller, 1998, from the lower Miocene (Karpatian) of Austria (Hyzny, 2016; Hyzny and Dulai, 2021). While any of the fossil metopograpsid species could serve as potential good candidates as vetted calibration points for crown Grapsidae, the fragmentary holotypes missing the fronto-orbital margins make their systematic placement less confident than those fossils referred to *P. hungaricus*.

The extinct genus *Miograpsus* Fleming, 1981, and its sole species *M. papaka* Fleming, 1981, is represented by a well-preserved female holotype and a ventrally exposed paratype from a “silty sandstone pebble, not in place” (*ex-situ*) (Fleming, 1981, p. 105), presumably from the lower Hurupi Formation, lower part of Tongaporutuan Stage (upper Miocene) of New Zealand. Although the genus *Miograpsus* has been placed within Grapsidae by several authors (e.g., Feldmann, 1993; Schweitzer et al., 2010), it seems to have closer affinities to genera within ‘Varuninae’ (now considered its own family, Varunidae) (e.g., Fleming, 1981; Karasawa and Kato, 2001; Webber et al., 2010). Since *Miograpsus* is 1) an extinct genus, 2) slightly younger that the fossil record of crown genera such as *Pachygrapsus* and *Metopograpsus*, and especially 3) may have putative varunid affinities, we refrain from using this taxon as calibration point for crown Grapsidae. ‘*Nautilograpsoides*’ *prior* Smirnov, 1929, a fossil species from the lower Miocene of the Caucasus, is currently envisioned as a juvenile form of the portunid genus *Liocarcinus* (Hyzný et al., 2022, and references therein).

*Pachygrapsus*, on the other hand, has a couple of fossil occurrences that can be assigned to this genus. Fragmentary cheliped remains of *Pachygrapsus* sp. are known from the upper Pleistocene of Jamaica (Morris, 1993; Luque et al., 2017), and *P. hungaricus* is known from several tens of specimens from the middle Miocene (Badenian) of Europe (see above). Since *Pachygrapsus* is an extant genus with a good Miocene fossil record, as represented by the type series and numerous additional specimens of *P. hungaricus*, we selected the latter as our calibration point for crown Grapsidae. The presence of *Pachygrapus* and *Metopograpsus* in the Miocene indicates that the most recent common ancestor of crown Grapsidae and all of its descendants must have a pre-Miocene origin, and likely rooted into the early Paleogene or Late Cretaceous.

## 13. Brachyura: Eubrachyura: Grapsoidea: Varunidae (crown)

### Fossil specimen

*Brachynotus corallinus* Beschin, Busulini, De Angeli & Tessier, 2007. Museo di Archeologia e Scienze Naturali “G. Zannato” di Montecchio Maggiore, Vicenza, northern Italy. Holotype MCZ 1794, a nearly complete dorsal carapace (Fig 3H).

### Phylogenetic justification

The overall carapace morphology indicates that the holotype of *Brachynotus corallinus* can be referred to crown Varunidae, which is fairly distantly related to Grapsidae, the eponym of the polyphyletic superfamily Grapsoidea (Chen et al., 2019; Lü et al., 2022; Tsang et al., 2022; Wolfe et al., 2022)

*Minimum age*. 47.8 Ma.

*Tip maximum age.* 56.0 Ma.

*Soft maximum age.* 100.9 Ma.

### Age justification

This coral-associated fossil was collected from lower Eocene (Ypresian) marine calcareous rocks, exposed at the Contrada Gecchelina of Monte di Malo in Vicenza, northern Italy.

The faunal association includes foraminifera, corals, calcareous algae, and molluscs. Foraminifera like *Nummulites* cf. *partschi*, *N. tauricus*, and *N. nitidus*, indicate a middle–upper Ypresian age (Beschin et al., 2007), and are stratigraphically correlated with the nearby ‘Rossi’ quarry (cava ‘Rossi’) (Beschin et al., 1998).

### Discussion

Other Paleogene fossils of crown Varuninae include *Brachynotus oligocenicus* De Angeli, Garassino and Ceccon, 2010b, a nicely preserved dorsal carapace from the early Oligocene of Vicenza, Italy (De Angeli et al., 2010b), confirming the presence of the genus and the family early in the Paleogene.

## 14. Brachyura: Eubrachyura: Grapsoidea: Gecarcinidae (crown)

### Fossil specimen

*Cardisoma guanhumi* Latreille, *in* Latreille, Le Peletier, Serville and Guérin, 1828. NCB – Naturalis, Leiden, The Netherlands, specimen NNM RGM 544 482, a large and mostly complete right chela preserving the dactylus and pollex.

### Phylogenetic justification

*Cardisoma guanhumi* is an extant gecarcinid species closely related to *Gecarcinus*, the type genus of the family Gecarcinidae. The phylogenetic position of Gecarcinidae is contentious with respect to other families within Grapsoidea. Recent nuclear and mitochondrial phylogenetic studies have recovered a non-monophyletic Grapsoidea, with gecarcinids closer to taxa within the families Sesarmidae and even Varunidae than to any taxon within Grapsidae (e.g., Chen et al., 2019; Liu et al., 2021; Lü et al., 2022; Tsang et al., 2022; Wolfe et al., 2022).

***Minimum age***. 0.0117 Ma.

*Tip maximum age.* 0.129 Ma.

*Soft maximum age.* 100.9 Ma.

### Age justification

This fossil was collected in the Port Morant Formation, parish of St. Thomas, east Port Morant Harbour, southeast Jamaica (Collins and Donovan, 1997, 2012). The Port Morant Formation, has been dated as Middle-Upper Pleistocene (132 ± 7 kyr to 125 ± 9 kyr, Sangamonian Stage) based on electron spin resonance dating on samples of the coral species *Solenastrea bournoni* and *Siderastrea radians* (Mitchell et al., 2000; Mitchell et al., 2001; James-Williamson and Mitchell, 2012). Given the age uncertainty of the fossil specimens from the Port Morant Formation with respect to the coral samples dated, we bracketed the soft maximum and minimum ages for the fossil occurrences of *C. guanhimu* between the base and the top of the Late Pleistocene (0.129– 0.0117 Ma), although we expect to family as a whole to be much older than Pleistocene, most likely originating in the Paleogene or possibly late Cretaceous (see in Wolfe et al., 2022).

Soft maximum age as for calibration 8.

### Discussion

The fossil record of gecarcinid land crabs is sparse and fragmentary, and largely represented by isolated claw remains (e.g., Luque, 2017) (Fig. 3I). To date, all of the known fossils assignable to *Cardisoma* are Quaternary in age, i.e., late Pleistocene (Collins and Donovan, 1997; Luque, 2017) or late Holocene (Luque, 2017; Luque et al., 2017; Luque et al., 2018), making the record from the upper Pleistocene of Jamaica the oldest fossil material referable to Gecarcinidae with confidence.

## 15. Brachyura: Eubrachyura: Grapsoidea: Sesarmidae (crown)

### Fossil specimens

Sesarmidae gen. et sp. indet., in Serrano-Sánchez et al. (2016). Museo de Paleontología “Eliseo Palacios Aguilera” (Secretaría de Medio Ambiente e Historia Natural), Calzada de los Hombres Ilustres s/n, Tuxtla Gutiérrez, Chiapas, Mexico, specimens with acronym IHNFG (Instituto de Historia Natural, Fósil Geográfico) IHNFG-4969, IHNFG-4970, IHNFG-4991, IHNFG-4992, and IHNFG-5555. A handful of specimens preserved in amber, mostly represented by complete dorsal and ventral carapaces with limbs attached (Serrano-Sánchez et al., 2016, figs 3, 4.1, 4.3–47; Luque et al., 2017, fig. 13L) (Fig. 3J).

### Phylogenetic justification

Despite the unclear generic and specific affinities of the fossil grapsoids in amber from the Miocene of Chiapas, Mexico, the fossils can be assigned with confidence to crown Sesarmidae based on the overall morphology of the carapace and limbs (Serrano-Sánchez et al., 2016; Luque et al., 2017).

*Minimum age*. 23.0 Ma.

*Tip maximum age.* 23.03 Ma.

*Soft maximum age.* 100.9 Ma.

### Age justification

These fossils were collected in rocks of the upper La Quinta Formation (Finca Carmitto Member) in Chiapas, México, dated as Lower Miocene (Aquitanian, 22.8 Ma, in Serrano-Sánchez et al., 2015) based on the biostratigraphy of corals, molluscs, microfossils, and Strontium isotopes (Vega et al., 2009; Serrano-Sánchez et al., 2015). Bernot et al. (2022) highlighted that the absence of radiometric ages for La Quinta Formation precludes a more precise constrain of the possible age of the crab-bearing amber samples, and propose some tentative minimum ages based on previously reported benthic foraminiferal data from the overlying Mazantic Shale (Solórzano Kraemer, 2007), while indicating a ^87^Sr/^86^Sr radiometric date of 23.0 Ma, roughly at the Oligocene–Miocene boundary, reported in Vega et al. (2009) (see Bernot et al. (2022) supplementary materials and reference therein). As such, given the uncertainty on the age of the crab-yielding amber deposits, we constrain the minimum age to the date reported by Serrano-Sánchez et al. (2015), and the tip maximum age to the base of the Aquitanian and thus the Miocene (23.03 Ma).

Soft maximum age as for calibration 8.

### Discussion

Sesarmid crabs trapped in amber are known from the lower Miocene (Aquitanian) of Simojovel, Chiapas, Mexico (Grimaldi, 1996; Boucot and Poinar Jr, 2010; Serrano-Sánchez et al., 2016; Luque et al., 2017). ‘*Sesarma*’ *paraensis* Beurlen, 1958, presumably a fossil sesarmid from the Miocene Pirabas Formation of Pará, Brazil, cannot be confirmed to be a sesarmid as the specimen is not illustrated in the original paper by Beurlen (1958), but only a schematic line drawing reconstruction (Beurlen, 1958, pl. IV, fig. 4). As such, as it stands today, this record cannot be reliably confirmed as a sesarmid. The only other sesarmid fossils known are cheliped remains from the upper Pleistocene of Jamaica previously described as a new species, *Sesarma primigenium* Collins, Mitchell and Donovan, 2009, but now considered a junior synonym of the extant species *Sesarma cookei* Hartnoll, 1971, to which the fossilized cheliped remains above mentioned belong (see discussion in Luque et al., 2017, p. 68, note 3).

## 16. Brachyura: Eubrachyura: Pseudothelphusoidea: Pseudothelphusidae (crown)

### Fossil specimen

Pseudothelphusidae gen. & sp. indet. (in Luque et al., 2019a). Invertebrate Paleontology collections, Florida Museum of Natural History. Specimen UF XXXXX, a complete dorsal and ventral carapace, chelipeds, eyes, and proximal podomeres of pereopods, in volume (Fig. 4A).

### Phylogenetic justification

The detailed preservation of diagnostic features in the fossil specimen, together with its stratigraphic and paleoenviromental context, e.g., continental leaf-rich level of the Pedro Miguel Formation, allow its placement with certainty within the freshwater group Pseutothelphusidae, and thus as part of the crown Pseudothelphusoidea (Luque et al., 2019a; Luque et al., under study).

*Minimum age*. 17.93Ma.

*Tip maximum age.* 18.18 Ma.

*Soft maximum age.* 100.9 Ma.

### Age justification

This fossil pseudothelphusid specimen comes *in-situ* from a leaf-rich level of tuffaceous mudstones of the Pedro Miguel Formation, Panama Canal, Panama. U/Pb absolute ages from detrital zircons collected *in-situ* in two tuffaceous conglomerates that bracket a leaf-rich horizon reported by Londoño et al. (2018) from which the fossil pseudothelphusid comes, yielded an estimated depositional age of 18.01±0.17 Ma. Previous Ar/Ar dating from the Pedro Miguel Formation in other parts of the Canal by Wegner et al. (2011) has suggested an age of 18.4 ± 1.07 Ma, and while details of the exact provenance of such samples within the formation are unclear (Montes et al., 2012; Farris et al., 2017), a corrected datum by MacFadden et al. (2014) suggests an age of 18.90±0.59 Ma.

Since the tuffaceous mudstone layer yielding the fossil leaves of Londoño et al. (2018) and the *in-situ* pseudothelphusid are bracketed by the same lower and upper tuffaceous conglomerates, both dated as 18.01±0.17 Ma, we assign the fossil a tip maximum age of 18.18 Ma, and a minimum age of 17.93 Ma, which are closer to the middle Burdigalian (late Early Miocene). As this fossil is clearly part of the crown Pseudothelphusidae (Luque et al., 2019a), the most recent common ancestor of all pseudothelphusids must have originated in the pre-Burdigalian, most likely the Paleogene.

Soft maximum age as for calibration 8.

### Discussion

There are only two extant superfamilies of freshwater crabs in the Neotropics: Trichodactylidae and Pseudothelphusidae. While the fossil record of Trichodactylidae is represented by numerous cheliped fragments (see below), the fossil record of Pseudothelphusidae is otherwise unknown. The new fossil represents not only the first and thus oldest know fossil of the family, but one of the most complete fossil freshwater crabs known to date (Luque et al., 2019a; Luque et al., under study).

## 17. Brachyura: Eubrachyura: Trichodactyloidea: Trichodactylidae (crown)

### Fossil specimen

Trichodactylidae genus and species indet. Museo de Historia Natural de la Universidad Nacional Mayor San Marcos de Lima (MUSM), Peru. Three claw fragments: CTA 47 (1 specimen) (Fig. 4B), and CTA 66 (two specimens), representing a small pollex (CTA 47), and a very small dactylus and a small pollex (CTA 66) (Klaus et al., 2017, fig. 3D–E).

### Phylogenetic justification

The preservation of diagnostic features on the fossilized dactyli and propodi and the teeth and interteeth on their cutting edges, together with their stratigraphic and continental limnic paleoenviromental contexts, allow their placement within the family Trichodactylidae (Klaus et al., 2017; Luque et al., 2017).

***Minimum age***. 40.94 Ma.

*Tip maximum age.* 43.44 Ma.

*Soft maximum age.* 100.9 Ma.

### Age justification

These fossils were collected in the lower member of the Pozo Formation, middle and upper middle Eocene (lower Barrancan, >41.6–40.94 Ma, 43.44 ± 2.5 Ma), Contamana area, Loreto, Peru. The ages are based on mammalian biochronology and ^40^Ar/^39^Ar radiometric age (Antoine et al., 2016; Klaus et al., 2017). The South American land mammal age (SALMA) known as Barrancan is largely equivalent to the uppermost global Lutetian and most of the Bartonian.

Soft maximum age as for calibration 8.

## 18. Brachyura: Eubrachyura: Epialtidae + Mithracidae (crown)

### Fossil specimen

**‘***Micippa*’ *antiqua* Beschin, De Angeli, and Checchi, 2001. Museo Civico “G. Zannato” di Montecchio Maggiore, Vicenza, northern Italy. Holotype I.G. 286477, a complete dorsal carapace (Fig. 4C).

### Phylogenetic justification

*Micippa* Leach, 1817, is an extant genus that has been recovered by several authors as not clustering together with other mithracids (see Windsor and Felder (2014) and references therein). Windsor and Felder (2014) included *Micippa* and *Stenocionops* Desmarest, 1823, within the family Mithracidae *sensu lato*, and Klompmaker et al. (2015) also included them within this family in their revision of the fossil record of Mithracidae (see discussion below). Alternatively, Poore and Ahyong (2023) include *Micippa* within Epialtidae. As such, we consider **‘***Micippa*’ *antiqua* as a calibration point for Epialtidae + Mithracidae (crown) (see in Wolfe et al., 2022). The fossil record of pre-Cenozoic majoid crabs is one of the most problematic for brachyurans due to the anatomical disparity seen across taxa and its often fragmentary nature, and it is in need of future thorough revisions.

***Minimum age***. 27.82 Ma.

*Tip maximum age.* 33.9 Ma.

*Soft maximum age.* 100.9 Ma.

### Age justification

The holotype of ‘*Micippa*’ *antiqua* was collected associated to shallow-water corals-bearing limestones from the Castelgomberto Formation (“Formazione di castelgomberto”) in Vicenza, northerm Italy (Beschin et al., 2001). Large foraminifera such as *Nummulites fichteli*, *N. vascus*, *Operculina complanata*, *Praerhapydionina delicata*, *Spirolina cylindracea*, *Peneroplis glynnjonesi*, and *Asterigerina rotula haeringensis* in the Castelgomberto Formation are indicative of the lower Oligocene (Rupelian, 33.9–27.82 Ma) biozones SB21–22A (Ungaro, 1978; Cahuzac and Poignant, 1997; Nebelsick et al., 2013).

Soft maximum age as for calibration 8.

### Discussion

Among the extant mithracid genera with a known fossil record, *Micippa* has the oldest putatively confirmed occurrences, as represented by ‘*M.*’ *antiqua* from the lower Oligocene of Italy. Dorsal carapaces of two other micippid species, i.e., *M. annamariae* Gatt and De Angeli, 2010, and *M. hungarica* (Lorenthey in Lorenthey & Beurlen, 1929), are known from the middle Miocene of Malta and the middle–upper Miocene of Poland, Hungary, and Austria, respectively. This, together with some Miocene and several Pliocene and Pleistocene fossils referable to Mithracidae s.s. from the Caribbean (Klompmaker et al., 2015; Luque et al., 2017), indicate that mithracids were already present and widespread during the late Paleogene–early Neogene.

‘*Stenocionops*’ *suwanneeana* Rathbun, 1935, from the upper Eocene fossil of Florida, USA, is a propodus that cannot be placed with confidence within any extant mithracid genus. “*Stenocionops*” *primus* Rathbun, 1935, presumably from the Upper Cretaceous (Santonian?) of Arkansas, USA, is a fragmented cheliped manus of unclear affinities. *Antarctomithrax thomsoni* Feldmann, 1994, from the Eocene of Antarctica, is an extinct monotypic genus described based on a partially preserved dorsal carapace that cannot be ascribed to Mithracidae with confidence.

## 19. Brachyura: Eubrachyura: Portunoidea: Eogeryonidae (stem)

### Fossil specimen

*Eogeryon elegius* Ossó, 2021. Museu de Geologia de Barcelona, holotype MGB 69151, a complete dorsal and ventral carapace in volume with cheliped associated (Ossó, 2016, 2021) (Fig. 4D).

### Phylogenetic justification

In a recent morphological phylogenetic study incorporating fossil and extant brachyurans, Eogeryonidae has been recovered as a stem Portunoidea, sister to a clade formed by the extant families Carcinidae, Geryonidae, and Portunidae, and all of them together forming the total group Portunoidea (Luque et al., 2021).

*Minimum age*. 93.9 Ma.

*Tip maximum age.* 97.0 Ma.

*Soft maximum age.* 100.9 Ma.

### Age justification

The type material of *Eogeryon elegius* comes from sandy limestones of the Villa de Vés Formation, exposed near the village of Condemios de Arriba, Northern Guadalajara Province, Spain. The presence of the ammonite *Vascoceras gamai* Choffat, 1898, in the Villa de Vés Formation represents the upper part of the upper Cenomanian (lowermost Upper Cretaceous, 95–93 Ma), and it is found below the *Spathites* (*Jeanrogericeras*) *subconciliatus* Zone, which marks the end of Cenomanian (Ossó, 2016).

Soft maximum age as for calibration 8.

### Discussion

The anatomical similarities of *Eogeryon* to portunoids such as *Geryon* suggest a potential proximity of Eogeryonidae to Geryonidae (Ossó, 2016, 2021). Several small dorsal carapaces of the eubrachyuran *Romualdocarcinus salesi* Prado and Luque *in* Prado et al., 2018, from the upper Lower Cretaceous (Aptian–Albian, ∼115–110 Ma) of Brazil, share some diagnostic features with *Eogeryon elegius* that suggest a plausible eogeryonid affinity, but due to their incompleteness, especially regarding the lack of ventral and cheliped material associated, a more precise phylogenetic placement of *R. salesi* is not possible at this time (Prado et al., 2018). *Eogeryon*, *Romualdocarcinus*, and other modern-looking crown eubrachyuran fossils, also share several general features with the remarkably well-preserved *Cretapsara athanata* Luque *in* Luque, Xing et al., 2021, from the lowermost Cenomanian (∼99 Ma) of Burma (Myanmar), but *Cretapsara* differs from them in having a bilobate rostrum and lacking orbital fissures (Luque et al., 2021).

## 20. Brachyura: Eubrachyura: Portunoidea: Geryonidae (crown)

### Fossil specimen

*Chaceon helmstedtense* (Bachmayer and Mundlos, 1968) (as *Coeloma* ?). Mundlos collection, geological-palaeontological collections of the Natural History Museum at Vienna, Austria. Holotype 1968/773/2, a complete dorsal carapace in volume with chelipeds and legs preserved (Fig. 4E).

### Phylogenetic justification

The fossil species *Chaceon helmstedtense* can be assigned to the extant genus *Chaceon* Manning and Holthuis, 1989, and thus the family Geryonidae, based on diagnostic features of the dorsal carapace outline, the orbitofrontal and anterolateral margins, the thoracic sternum, and the chelipeds. Molecular and morphological phylogenetic studies have recovered *Chaceon* and Geryonidae as nested within crown Portunoidea (e.g., Karasawa et al., 2008; Spiridonov et al., 2014; Evans, 2018; Luque et al., 2021; Wolfe et al., 2022).

***Minimum age***. 27.82 Ma.

*Tip maximum age.* 33.9 Ma.

*Soft maximum age.* 97.0 Ma.

### Age justification

The type locality of *Chaceon helmstedtense* is a marl/clay horizon (Mergel/Ton-Horizont, crab zone K I) in Helmstedt near Braunschweig, Lower Saxony, Northwestern Germany, considered lower Oligocene in age (Rupelian, 33.9–27.82 Ma). Associated molluscan fauna in the marl/clay beds includes “*Pecten corneus*, *Isocardia multicostata*, *Cardita* latesulcata, *Ostrea ventilabrum*, and *Ostrea queteleti*“ (Bachmayer and Mundlos, 1968). As crown group members of Portunoidea cannot be older than their common ancestor shared with the stem group fossil *Eogeryon* (Ossó, 2016), a soft maximum age is taken from the maximum uncertainty on the age of the Villa de Vés Formation, at 97.0 Ma.

### Discussion

Besides *Chaceon helmstedtense*, another occurrence of *Chaceon* in the Paleogene is *C. heimertingensis* (Bachmayer and Wagner, 1957), from the uppermost Oligocene (Chattian) of Austria. *Chaceon peruvianus* (D’Orbigny, 1842, as *Portunus*) an iconic Miocene fossil crab from southern South America—mainly Argentina, but also Chile, Peru, and Uruguay (e.g., Casadío et al., 2005; Luque et al., 2017; Perea et al., 2020, and references therein). This species has been reported from the Estancia 25 de Mayo Formation (previously known as Centinela Formation) in Tierra del Fuego, Argentina, whose lower portion has been dated as upper Oligocene–Lower Miocene (Chattian–Aquitanian, 27.28–20.44 Ma) based on ^87^Sr/8^6^Sr radioisotopic dating ages between 25.5–21.5 Ma from a valve of the oyster *Crassostrea*? *hatcheri* (Ortmann), collected below a tuff bed near the Oligocene–Miocene (Chattian–Aquitanian) boundary (Casadío et al., 2000; Casadio et al., 2001; Guerstein et al., 2004), and the palynomorphs *Operculodinium israelianum*, *Spiniferites ramosus*, *Lingulodinium machaerophorum*, *Reticulatosphaera actinocoronata*, *Nematosphaeropsis* cf. *N. lativittata*, *Baumannipolis variapertatus*, *Mutisiapollis viteauensis*, *Nothofagidites* spp., and *Podocarpidites* spp. (Guerstein et al., 2004). As such, the most recent common ancestor of Geryonidae and all of its descendants must have originated in the pre-Oligocene.

## 21. Brachyura: Eubrachyura: Portunoidea: Portunidae: Thalamitinae (crown)

### Fossil specimen

*Lessinithalamita gioiae* De Angeli and Ceccon, 2015. Museo Civico di Valdagno, Italy. Holotype MCV14/15, a fairly complete dorsal carapace (Fig. 4F).

### Phylogenetic justification

The extinct, monotypic genus *Lessinithalamita* De Angeli and Ceccon, 2015, shows remarkable affinities with the extant genera *Thalamita* Latreille, 1829, and *Thranita* Evans, 2018, justifying its inclusion within the subfamily Thalaminitae, which is nested within the crown group Portunidae (Evans, 2018).

*Minimum age*. 47.8 Ma.

*Tip maximum age.* 53.3 Ma.

*Soft maximum age.* 97 Ma.

### Age justification

The specimen studied comes from compact calcarenites rich in coralline algae (e.g., *Lithothamnium bolcensis*) and calcareous nannofossils rocks cropping out near Monte Magrè, Vicenza, Italy, and dated approximately as lower Eocene (middle and upper Ypresian, ∼53.3–47.8 Ma) (Beccaro, 2003, in De Angeli and Ceccon, 2012).

Soft maximum age as for calibration 20.

### Discussion

Besides *Lessinithalamita gioiae*, there are two other species of Eocene thalamitine crabs belonging to the extinct genus *Eocharybdis* Beschin, Busulini, De Angeli & Tessier, 2002, i.e., *E. rugosa* Beschin, Busulini, Tessier & Zorzin, 2016a, from the lower Eocene (Ypresian) of Monte di Malo, Verona, NE Italy, and *E. cristata* Beschin, Busulini, De Angeli & Tessier, 2002, from the middle Eocene (Lutetian) Cava “Main” di Arzignano, Vicenza, Italy. While both of these species are suitable candidates for calibration points for the subfamily Thalamitinae, the completeness of the type material of *L. gioiae* and its similarities to *Thalamita* and *Thranita*, support a more reliable affiliation to the crown Thalamitinae.

## 22. Brachyura: Eubrachyura: Portunoidea: Polybiidae (crown)

### Fossil specimen

*Liocarcinus heintzi* Schweitzer and Feldmann, 2010b. Muséum national d’Histoire naturelle, Paris. Holotype MNHN R03778, a well-preserved dorsal carapace (Fig. 4G).

### Phylogenetic justification

*Liocarcinus* Stimpson, 1858, is an extant portunoid genus nested within crown Polybiidae (Evans, 2018). The overall sub-hexagonal carapace of *L. heintzi*, slightly wider than long, with four anterolateral spines, the concave posterolateral margins, a convex posterior margin, the orbits directed forward, and marked dorsal regions, seem to conform with the forms seen among species of *Liocarcinus* (Schweitzer et al., 2021c).

***Minimum age***. 27.82 Ma.

*Tip maximum age.* 33.9 Ma.

*Soft maximum age.* 97 Ma.

### Age justification

The studied specimen comes from the carbonatitic Calcaire à Astéries Formation, cropping out near Monségur, Gironde, Western Aquitaine, France (Schweitzer and Feldmann, 2010b). The Calcaire à Astéries Formation has been dated as lower Oligocene (Rupelian, 33.9–27.82 Ma), based on the recognition of the foraminifera biozones SBZ 21 and 22A, containing *Nummulites fichteli*, *N. intermedius N. vascus*, *Spiroclypeus*, *Operculina*, *Halkyardia* spp., *Neorotalia* spp., *Peneroplis*, and *Arenagula* and the ostreid *Crassostrea longirostris* (Cahuzac and Londeix, 2012; Sztrákos and Steurbaut, 2017).

Soft maximum age as for calibration 20.

### Discussion

Besides *Liocarcinus heintzi*, there are several other occurrences of polybiid and polybiid-like genera and species in the Paleogene. *Archaeogeryon corsolini* (Casadío, De Angeli, Feldmann, Garassino, Hetler, Parras and Schweitzer, 2004, as *Proterocarcinus*), and *A. lophos* (Feldmann, Casadío, Chirino-Gálvez, and Aguirre-Urreta, 1995), are known from the middle Oligocene and Paleocene (Danian, 66.0–61.6 Ma) of Argentina, respectively (Feldmann et al., 1995; Casadío et al., 2004; Casadío et al., 2005; Feldmann et al., 2011b). The latter species, represented by the holotype and sole specimen, is moderately preserved and shows some dorsal and sternal features that suggest affinity with portunids in general, but the fronto-orbital margin and part of the anterolateral margins are eroded, precluding a detailed comparison with polybiids in general.

Moreover, the exact phylogenetic position of *Archaeogeryon* Colosi, 1924, among portunoids, as well as that of the also extinct genus *Gecchelicarcinus* Beschin, Busulini, De Angeli, and Tessier, 2007, known from two species from the Ypresian of Italy, is unclear. Both taxa could represent extinct polybiid genera, but they cannot be assigned to the crown group with certainty.

The presumably polybiid monotypic extinct genera *Boschettia* Busulini, Tessier, Beschin, and De Angeli, 2003, from the lower and middle Eocene (Ypresian to Lutetian, 56.0–41.2 Ma) of Italy, and *Falsiportunites* Collins and Jakobsen, 2003, from the middle Eocene (?Lutetian, 47.8–41.2 Ma) of Denmark, show a combination of anatomical features on their fronto-orbital margins, anterolateral margins, and dorsal carapaces that differ from those seen among polybiids and thus cannot be accommodated satisfactorily among the crown group either.

Among the extant genera with known fossil records, *Liocarcinus oligocaenicus* (Pauca, 1929, as *Portunus*), and *L. atropatanus* (Aslanova and Dschafarova, 1975) are known from the lower and upper Oligocene (Rupelian–Chattian, 33.9–23.3 Ma) of Romania and Azerbaijan, respectively (Schweitzer et al., 2009; Beschin et al., 2016b; Hyzny, 2016), which confirm the presence of the genus during Oligocene times. *Liocarcinus priscus* Beschin, De Angeli, Checci, and Zarantonello, 2016b, from the lower Eocene (lower Lutetian, 47.8–41.2 Ma), may represent the oldest occurrence of the genus and thus a more approximate soft maximum age of calibration for the polybiid node, but due to its remarkable differences with the type species of the genus *Liocarcinus,* i.e., *L. holsatus* (Fabricius, 1798), to which *L. heintzi* is anatomically closer, we favor the younger fossil for a calibration point. Regardless of which of these fossils is considered the oldest reliable record of the family, the clear occurrence of several Paleogene genera and species attributable to Polybiidae indicate that the most recent common ancestor of the family and all of its descendants originated most likely during the early Paleogene (Wolfe et al., 2022), or even the Late Cretaceous.

## 23. Brachyura: Eubrachyura: Parthenopoidea: Parthenopidae+Dairoididae (crown)

### Fossil specimen

*Aragolambrus collinsi* Ferratges, Zamora, and Aurell, 2019. Palaeontological collection of the Museo de Ciencias Naturales de la Universidad de Zaragoza, Spain. Holotype MPZ-2019/211, a nearly complete dorsal carapace with chelipeds associated (Fig. 4H).

### Phylogenetic justification

In a recent molecular phylogenetic study, Wolfe et al. (2022) recovered *Dairoides kusei* as sister to a monophyletic Parthenopidae, whereas Ferratges et al. (2023), based on a morphological phylogenetic study of Parthenopoidea, recovered the extinct monotypic genus *Aragolambrus* Ferratges, Zamora, and Aurell, 2019, in close proximity to the also extinct genus *Phrynolambrus* Bittner, 1893, and the extant genus *Dairoides* Stebbing, 1920, all three forming a well-supported monophyletic subfamily Dairoidinae sister to Daldorfiinae and nested within a paraphyletic Parthenopidae (Ferratges et al., 2023). Future comprehensive molecular phylogenetic studies with larger taxon coverage across multiple parthenopoid genera will shed light on the internal relationships among these groups.

***Minimum age***. 53.02 Ma.

*Tip maximum age.* 57.1 Ma.

*Soft maximum age.* 100.9 Ma.

### Age justification

*Aragolambrus collinsi* comes from the coral-algal Reef Limestones interval (middle member) of the Serraduy Formation, in the Barranco de Ramals outcrop near Puebla de Roda and Serraduy, Huesca province, Spain (Ferratges et al., 2019). The Reef Limestones interval develops over the underlying Alveolina Limestones interval and the overlying marls of the Riguala Member (Serra-Kiel et al., 1994; Pujalte et al., 2009). Serra-Kiel et al. (1994) assigns an age of lower Ilerdian to the Alveolina Limestones interval, which is located within the magnetostratigraphic Chron C24r (57.101–53.9 Ma, Francescone et al., 2019; Ogg, 2020), and contains the macroforaminifera *Alveolina cucumiformis, A. ellipsoidalis*, and *Nummulites fraasi*, from the NP9, BB1, and P6a Biozones. For the marls of the Riguala Member, the same authors assigns an age of middle Ilerdian, roughly located between the top of the magnetostratigraphic Chron C24r and Chron C24n.1r (∼53.9–53.02 Ma, Francescone et al., 2019; Ogg, 2020), and contains the macroforaminifera *Alveolina moussoulensis*, *A. corbarica*, *Nummulites robustifrons,* and *N. exilis*, from the BB1–BB2, NP9–NP10, and P6a-P6b Biozones (Serra-Kiel et al., 1994). Due to the lateral facial discontinuity of the Reef Limestones interval, which is bounded at the bottom by the Alveolina Limestones and at the top by the Riguala Member, we bracket the tip maximun and minimum ages of *A. collinsi* between 57.1–53.02 Ma, with a possible age of the fossil itself closer to 53.5 Ma (F. Ferratges, pers. comm. to J. Luque, Feb. 2023).

Soft maximum age as for calibration 8.

### Discussion

*Aragolambrus* and *Phrynolambrus* are two of the oldest fossils of crown Parthenopidae known, sharing with *Dairoides* the presence of two to three anterolateral spines, an ornamented epistomia with one or more rows of tubercles, and the pterygostome with grooves, which are synapomorphies that unite them among parthenopoids under the subfamily Dairoidinae (Ferratges et al., 2023).

The extinct genera *Eogarthambrus* De Angeli, Garassino, and Alberti, De Angeli et al., 2010a, and *Mesolambrus* Müller and Collins, 1991, both known from the Eocene (middle Ypresian to Priabonian, ∼52.0–33.9 Ma) of Italy, were recovered by Ferratges et al. (2023) outside the crown Parthenopoidea. The monotypic extinct genus *Braggilambrus* De Angeli and Caporiondo, 2016, from the Ypresian of Italy, seems to fit within the general parthenopoidean body form, but due to its incompleteness and therefore lack of key diagnostic characters, its affinities at the family and subfamily levels is still unclear.

The presence of several crown parthenopid forms across the early to late Eocene indicate that the most recent common ancestor of Parthenopoidea and all of its descendants must have Paleocene or even Late Cretaceous origins.

## 24. Brachyura: Eubrachyura: Calappoidea: Calappidae (crown)

### Fossil specimen

*Calappa zinsmeisteri* Feldmann and Wilson, 1988. Smithsonian Institution, National Museum of Natura History (NMHN), USA. Holotype USNM 404877, claws only (Fig. 4I,J).

### Phylogenetic justification

*Calappa* Weber, 1795, is the type genus of the family Calappidae De Haan, 1833, and thus phylogenetically nested within the crown Calappoidea (Lu et al., 2020).

*Minimum age*. 33.9 Ma.

*Tip maximum age.* 41.2 Ma.

*Soft maximum age.* 100.9 Ma.

### Age justification

The type material of *Calappa zinsmeisteri* comes from the uppermost La Meseta Formation on Seymour Island, Antarctica, Locality 14, of Feldmann and Wilson (1988). This locality corresponds to the Telms VI–VII of Sadler (1988), which have been dated as middle to upper Eocene (Bartonian–Priabonian, ∼42.0–34.0 Ma) based on ^87^Sr/^86^Sr radiometric dating ages from *Cucullaea* shells from Telm VII between 46.13 and 34.69 Ma (Dutton et al., 2002), and the occurrences of the neogastropods *Prosipho lawsi*, *P. lamesetaensis*, *Austroficopsis meridionalis*, *Microfulgur byrdi*, *Fusinus*? *suraknisos*, *Eupleura suroabdita*, ?*Adelomelon suropsilos*, *Zemacies finlayi*, and *Aforia canalomos* (Crame et al., 2014).

Soft maximum age as for calibration 8.

### Discussion

Among the extant calappid genera, only *Calappa* and *Mursia* Leach, *in* Desmarest, 1823, have recognized Eocene fossils, to our knowledge. Four species of *Calappa*—including *C. zinsmeisteri*— are known from the middle and upper Eocene of USA, Antarctica, and presumably Venezuela, and are constituted largely by cheliped (propodi) remains with diagnostic anatomical details that can be assigned with certainty to crown Calappidae, and most likely to *Calappa* (Ross et al., 1964; Feldmann and Wilson, 1988; Luque et al., 2017). One species of *Mursia* Leach, *in* Desmarest, 1823, i.e., *Mursia aspina* Schweitzer and Feldmann, 2000a, from the Late Eocene of Washington, USA, is represented by a handful of dorsal carapaces that strongly resemble *Mursia*, although they lack some of the diagnostic features of the genus such as developed lateral spines. While we considered the cheliped remains of *Calappa zinsmeisteri* as our calibration point given their putative affinities to the type genus of the family, *M. aspina*, which is represented by dorsal carapace material of similar age as *C. zinsmeisteri*, could also be a good candidate to calibrate the node for crown Calappidae.

Among the extinct calappid genera, *Calappilia* A. Milne-Edwards in de Bouillé, 1873, bears conspicuous anatomical similitudes with *Calappa*, and has over 20 species known, of which more that 15 are Eocene in age (e.g., Williams and Child, 1988; Schweitzer et al., 2010; Rumsey et al., 2016; Luque et al., 2017; Feldmann et al., 2019, and references therein). There are also several extinct genera known exclusively from the Eocene, such as the monotypic genera *Paracorallomursia* Beschin, Busulini, Tessier & Zorzin, 2016a, and *Pseudocorallomursia* Beschin, Busulini, Tessier & Zorzin, 2016a, from the Ypresian of Italy, *Carinocalappa* Beschin, Busulini, and Tessier *in* Beschin et al., 2018, from the Priabonian of Italy, and *Tavernolesia* Artal and Onetti, 2017, from the Lutetian of Spain. *Corallomursia* De Angeli and Ceccon, 2014, comprises two species, both from the Ypresian of Italy (Beschin et al., 2015). All of these extinct genera seem to fit well within the total group Calappidae, but their precise affinities with crown calappids remain unclear.

Calappid crabs were already diverse, speciose, and widespread by the Eocene, indicating that the most recent common ancestor of Calappidae and all of its descendants must have originated in the earliest Paleogene (Paleocene) or the Cretaceous.

## 25. Brachyura: Eubrachyura: Cancroidea: Cancridae (crown)

### Fossil specimen

*Anatolikos undecimspinosus* Schweitzer, Feldmann, González-Barba, and Ćosović, 2006. Museo de Historia Natural, Universidad Autónoma de Baja California Sur, La Paz, Baja California Sur, México (MHN-UABCS). Holotype MHN-UABCS/Ba12-9, a partially preserved dorsal left carapace (Fig. 4K).

### Phylogenetic justification

The extant genus *Anatolikos* Schweitzer and Feldmann, 2000b, can be placed with confidence within the family Cancridae (Schweitzer et al., 2006; Schram and Ng, 2012; Wolfe et al., 2022). The diagnostic shape and arrangement of the eleven anterolateral spines seen in the extant genus *Anatolikos* and thus in *A. undecimspinosus*, together with the presence of a nearly straight rostrum with “five coalesced spines separated by fissures” (Schweitzer and Feldmann, 2000b) are unique traits not shared with other genera within the family (Schweitzer et al., 2006; Poore and Ahyong, 2023).

*Minimum age*. 41.2 Ma.

*Tip maximum age.* 56.0 Ma.

*Soft maximum age.* 100.9 Ma.

### Age justification

The type material of *Anatolikos undecimspinosus* comes from middle Eocene rocks of the Bateque Formation, cropping out at locality Waypoint 29 of Schweitzer et al. (2006), northwest of La Paz, near the village of San Ignacio, Baja California Sur, Mexico. The age of the Bateque Formation has been constrained between the early Eocene and the late middle Eocene (56.0–41.2 Ma) based on foraminiferal, nannoplankton, and diatom assemblages, more than 150 macroinvertebrates, and ostracods (Mina-Uhink, 1957; Sorensen, 1982; McLean et al., 1987; Carreño and Cronin, 1993; Morales-Ortega et al., 2016).

Soft maximum age as for calibration 8.

### Discussion

Among the cancrids with known fossil record, only the Cancrinae genera *Anatolikos* and presumably *Cancer* Linnaeus, 1758, have Eocene occurrences. However, *Cancer gabbi* Rathbun, 1926, from the Eocene of California, is represented by claw fragments only that cannot be confirmed to belong to the group. *Cancer meticuriensis* Thurmann, 1853, from the Oligocene and presumably from the lower Eocene of France, is too incomplete, with only the sternum and the underside of the front legs known, which A. Milne-Edwards (1865) considered to most likely belong to a portunid. At least six species (*C. aldenardensis*, *C. archiaci*, *C. flandricus*, *C. pratti*, 1850, *C. rotnacensis*, and *C. villabersiani*) from the Ypresian–Lutetian of Europe, are all *nomina nuda*, many of which have never been described or figured, and therefore their generic affinities are dubious (Van Straelen, 1920; Nations, 1975). *Cancer santosi* (Rathbun, 1937, as *Lobocarcinus*?) from the upper Eocene of Panama, is a poorly preserved right dorsal carapace.

Three extinct monotypic genera with potential Carcininae affinities have also early to middle Eocene occurrences, i.e., *Santeecarcinus* Blow and Manning, 1996, and *Sarahcarcinus* Blow and Manning, 1996, both from USA, and *Notocarcinus* Schweitzer and Feldmann, 2000b, from Argentina. However, given that these three genera are extinct only, we consider them suboptimal candidates for the calibration of the Cancridae node, given the presence of fossil taxa assigned to extant genera, e.g., *Anatolikos undecimspinosus*, in rocks of similar ages.

Several other extinct cancroid genera currently placed within the subfamily Lobocarcininae Beurlen, 1930 (e.g., De Grave et al., 2009; Schweitzer et al., 2010), also have their oldest or only records in the Eocene. These include *Miocyclus* Müller, 1978, from Bulgaria, *Nicoliscarcinus* Beschin, Busulini, Tessier & Zorzin, 2016a, and *Ramacarcinus* De Angeli & Ceccon, 2017, from Italy, and *Tasadia* Müller, *in* Janssen and Muller, 1984, from Belgium. In addition, two out of the three species of the extinct genus *Ceronnectes* De Angeli and Beschin, 1998, and several species of *Lobocarcinus* Reuss, 1857, also occur in the middle Eocene (e.g., Anderson and Feldmann, 1995; De Angeli and Beschin, 1998; Feldmann et al., 1998; Schweitzer et al., 2010; Beschin et al., 2016a).

## 26. Brachyura: Eubrachyura: Dorippoidea (crown)

### Fossil specimen

*Bartethusa hepatica* Quayle and Collins, 1981. Palaeontological Department, London’s Natural History Museum (formerly British Museum (Natural History)), England, holotype BM In.61704, a well-preserved dorsal carapace (Fig. 5A).

**Figure 4.**
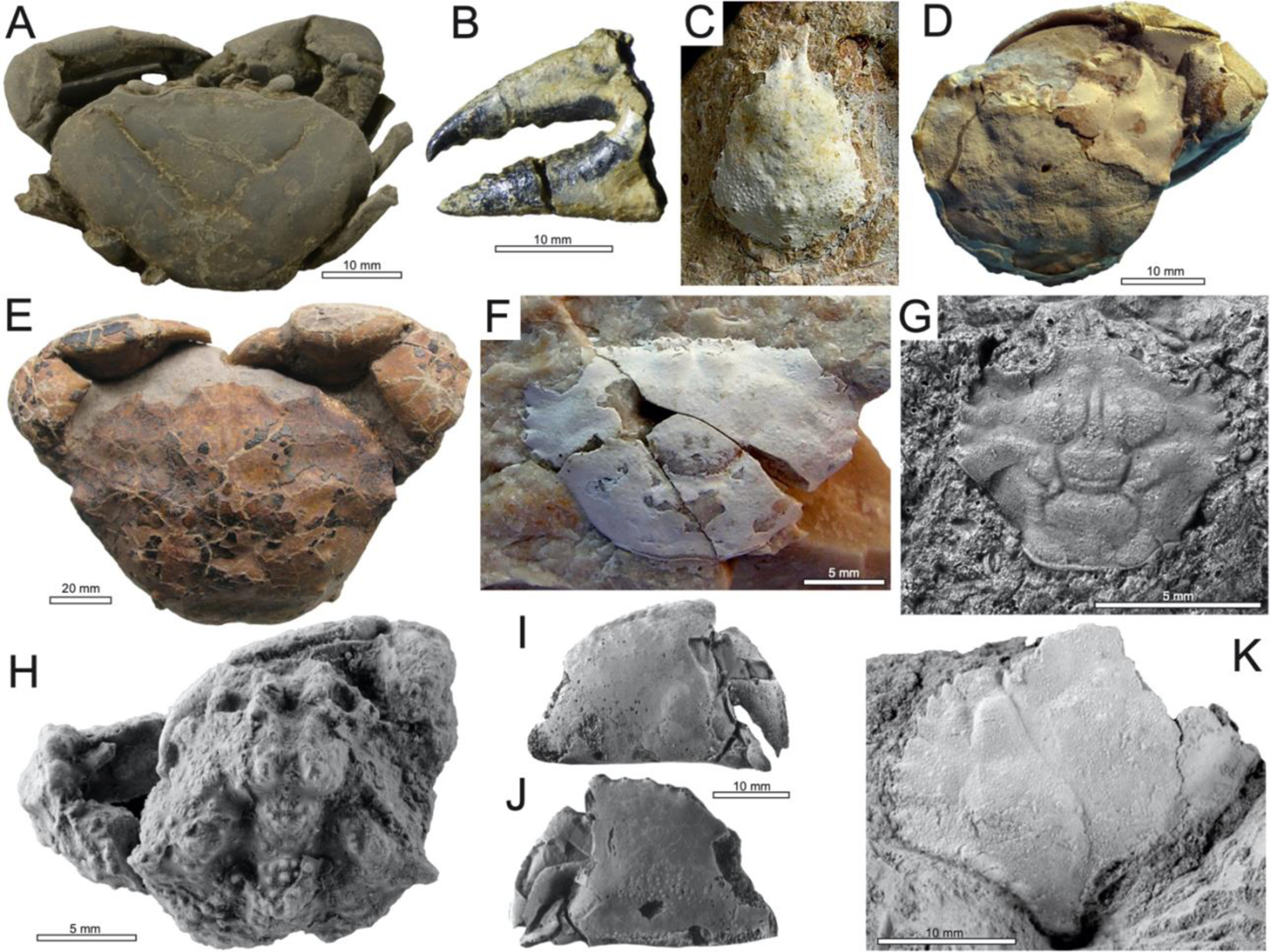
Crown Eubrachyura, cont. **A**: Pseudothelphusioidea: Pseudothelphusidae indet., specimen XXXX, Early Miocene (Burdigalian), Panama Canal, Panama (Luque et al., 2019a). **B**: Trichodactyloidea: Trichodactylidae indet., specimen CTA 47, Eocene (Barrancan), Contamana, Peru. **C**: Majoidea: Epialtidae + Mithracidae: *Micippa antiqua* Beschin, De Angeli, and Checchi, 2001, holotype I.G. 286477, lower Oligocene (Rupelian), Vicenza, Italy. **D–G**: Portunoidea; **D**: Eogeryonidae: E*ogeryon elegius* Ossó, 2021, holotype MGB 69151, lower Upper Cretaceous (Cenomanian), Northern Guadalajara Province, Spain (Ossó, 2016); **E**: Geryonidae: *Chaceon helmstedtense* (Bachmayer and Mundlos, 1968) (as *Coeloma*?), holotype Holotype 1968/773/2, lower Oligocene (Rupelian), Northwestern Germany; **F**: Portunidae: Thalamitinae: *Lessinithalamita gioiae* De Angeli and Ceccon, 2015, holotype MCV14/15, lower Eocene (Ypresian), Vicenza, Italy; **G**: Polybiidae: *Liocarcinus heintzi* Schweitzer and Feldmann, 2010b, holotype MNHN R03778, lower Oligocene (Rupelian), Western Aquitaine, France. **H**: Parthenopoidea: Parthenopidae + Dairoididae: *Aragolambrus collinsi* Ferratges, Zamora, and Aurell, 2019, holotype MPZ-2019/211, lower Eocene (Ypresian–Ilerdian), Huesca province, Spain. **I**: Calappoidea: Calappidae: *Calappa zinsmeisteri* Feldmann and Wilson, 1988, holotype USNM 404877, upper Eocene, Seymour Island, Antarctica. **J**: Cancroidea: Cancridae: *Anatolikos undecimspinosus* Schweitzer, Feldmann, González-Barba, and Ćosović, 2006, holotype MHN-UABCS/Ba12-9, middle Eocene, Baja California Sur, Mexico. Photos by: Javier Luque (A), Pierre Olivier Antoine (B), Antonio De Angeli (C and F), Àlex Ossó (D), Barry van Bakel (E), Rodney Feldmann (G, I–K), and Fernando Ferratges (H).

**Figure 5.**
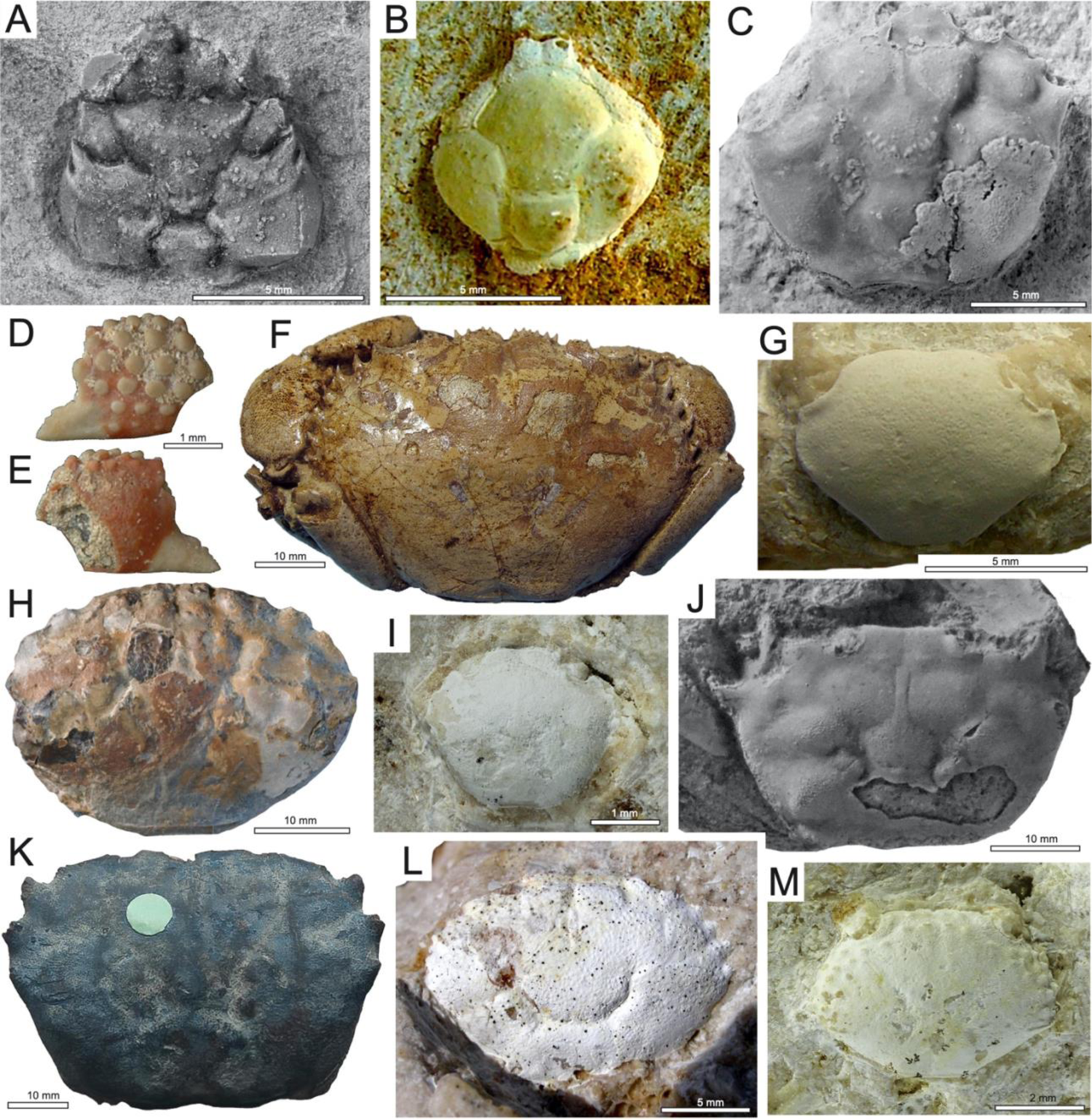
Crown Eubrachyura, cont. **A**: Dorippoidea: *Bartethusa hepatica* Quayle and Collins, 1981, holotype BM In.61704, middle Eocene (Bartonian), Isle of Wight, England. **B**: Leucosioidea: Leucosiidae: *Typilobus alponensis* Beschin, De Angeli, and Zorzin, 2009, holotype IGVR78539, lower Eocene (Ypresian), Verona, Italy. **C:** Goneplacoidea: Euryplacidae: *Chirinocarcinus wichmanni* (Feldmann, Casadío, Chirino-Galvez, and Aguirre-Urreta, 1995) (as *Glyphithyreus*), holotype GHUNLPam 7015, lowermost Paleocene (lower Danian), Neuquén Province, Argentina. **D–F**: Eriphioidea; **D–E**: Eriphiidae: *Eriphia verrucosa* (Forskal, 1775) (as *Cancer*), in Betancort et al. (2014), specimen number indet., upper Miocene, Islas Canarias, Spain. **F**: *Eriphia cocchi* Ristori, 1886, specimen C-021-2.1, lower Pliocene (Zanclean–Piacentian), Italy. **G**: Trapezioidea: Trapeziidae: *Archaeotetra lessinea* De Angeli and Ceccon, 2013, holotype MCV 12/05-I.G.360311, lower Eocene (Ypresian), Monte Magré, Lessini Mounts, Italy. **H**: Eriphioidea: Oziidae: *Ozius collinsi* Karasawa, 1992, holotype MFM39001, lowermost Middle Miocene, Okayama Prefecture, Japan, Japan. **I**: Pilumnoidea: Pilumnidae: *Glabropilumnus trispinosus* Beschin, Busulini, and Tessier, *in* Beschin et al., 2016a, holotype VR 94508, lower Eocene (Ypresian), Verona, Italy. **J**: Goneplacoidea: Goneplacidae: *Carcinoplax temikoensis* Feldmann and Maxwell, 1990, holotype AR 1943, upper Eocene (Kaiatan or Runangan), Westland, New Zealand. **K**: Xanthoidea: Pseudorhombilidae: *Pseudorhombila patagonica* Glaessner, 1933, holotype In. 28031, Miocene indet., Santa Cruz Province, Argentina. **L**: Xanthoidea: Panopeidae: *Panopeus incisus* Beschin, Busulini, De Angeli, and Tessier, 2007, holotype MCZ 2009, lower Eocene, Vicenza, Italy. **M**: Xanthoidea: Xanthidae: *Phlyctenodes edwardsi* Beschin, Busulini, Tessier, and Zorzin, 2016a, holotype VR 94284, lower Eocene (Ypresian), Verona, Italy. Photos by: Richard Howard (A, K), Antonio De Angeli (B, G), Rodney Feldmann (C, J), Juan Francisco Betancort (D, E), Àlex Ossó (F), Hiroaki Karasawa (H), Alessandra Busulini (I, L, M).

### Phylogenetic justification

*Bartethusa hepatica* has an overall carapace outline, fronto-orbital construction, and dorsal carapace region configuration that matches the range of forms seen across crown dorippoids. While *Bartethusa hepatica* could be allied to either Ethusidae Guinot, 1977 (e.g., Quayle and Collins, 1981; Van Bakel et al., 2021) or Dorippidae MacLeay, 1838a (e.g., Luque, 2015b; Schweitzer et al., 2021a; Guinot, 2023), in either case it will still be nested within the crown Dorippoidea MacLeay, 1838a.

***Minimum age***. 37.71 Ma.

*Tip maximum age.* 41.2 Ma.

*Soft maximum age.* 100.9 Ma.

### Age justification

The type material of *Bartethusa hepatica* was collected in rocks of the Horizon A3, Barton Beds, Christchurch Bay, Isle of Wight, England (Quayle and Collins, 1981), which have been dated as middle Eocene (Bartonian, 41.2–37.71 Ma) based on larger and smaller foraminifera, calcareous nannofossils, palynomorphs, and paleomagnetism (see Cotton et al., 2021, and references therein). Tsang et al. (2014) also used *B. hepatica* as their calibration point, but they assigned it an age of lower Eocene (Ypresian, 56.0–47.8 Ma). However, the available stratigraphic information indicates that the correct age of the known fossils of *B. hepatica* is middle Eocene (Bartonian, 41.2– 37.71 Ma) and not Ypresian, based on larger and smaller foraminifera, calcareous nannofossils, palynomorphs, and paleomagnetism (see Cotton et al., 2021, and references therein).

Soft maximum age as for calibration 8.

### Discussion

Dorippoid crabs—extant and fossil—are among the most puzzling of the eubrachyurans in terms of their phylogenetic affinities, given their unique combination of plesiomorphic and apomorphic characters and their relatively low representation in previous molecular phylogenies compared to most other crabs. Based on morphology alone, dorippoids have been typically regarded as some of the oldest crown eubrachyurans, and thus one of the earliest splitting branches in the eubrachyuran tree, with their earliest known representatives, e.g., Telamonocarcinidae Larghi, 2004, and potentially Tepexicarcinidae Luque, 2015b, already present in the Early Cretaceous (Luque, 2015b; Luque et al., 2021; Van Bakel et al., 2021). Although their overall anatomy is strongly reminiscent of the array of body forms across modern dorippoids in terms of their dorsal carapaces, thoracic sternum, and shape and size of their pereopods (P2–P5), they cannot be assigned to any of the two crown families recognized today, i.e., Dorippidae and Ethusidae. Some of these other extinct families, however, may represent convergent morphology, as dorippoids consistently nest well within heterotremes in molecular phylogenies (Tsang et al., 2014; Wolfe et al., 2022).

Among the extinct post-Cretaceous dorippoid forms, *Goniochele* Bell, 1858, presumably belonging to its own family Goniochelidae Schweitzer and Feldmann, 2011, has a confirmed fossil record restricted to two Eocene species: *G. angulata* Bell, 1858, from the Ypresian of England, and *G. madseni* Collins and Jakobsen, 2003, from the Ypresian–Lutetian of Denmark. However, the few similarities between Goniochelidae and some members of Dorippoidea s.s. might be convergent and not reflecting shared ancestry (Guinot, 2023). Rathbun (1918 [1919]), based on an isolated dactylus from the Early Miocene (not Oligocene, as Rathbun suggested) Culebra Formation in the Panama Canal, erected a third species of *Goniochele*, *G. armata* Rathbun, 1918 [1919], that currently cannot be ascribed to this genus with certainty (see Luque et al., 2017). Another extinct taxon, *Archaeocypoda veronensis* Secrétan, 1975, known from the lower Eocene (upper Ypresian) of Italy (Pasini et al., 2019), has been presumably assigned to Dorippidae based on the sub-oval carapace outline, the broad front and continuous supraorbital margin, the smooth anterolateral and posterolateral margins, the inflated protogastric and branchial regions, the smooth antero- and posterolateral margins (Casadío et al., 2005; Pasini et al., 2019; Schweitzer et al., 2021a). While these features match some of the diagnostic characters of dorippids, the possession of at least three pairs of well-developed pereopods (i.e., P2–P4) dramatically differs from the apomorphic dorippoid possession of two large pereopods (P2–P3) and two reduced, subchelate, and sub-dorsally carried pereopods (P4–P5), which is a feature seen across crown dorippoids and even stem dorippoids from the Cretaceous (e.g., Larghi, 2004; Casadío et al., 2005; Luque, 2015b; Guinot et al., 2019; Guinot, 2023). As such, while *A. veronensis* might (or might not) be related to crown Dorippoidea, its systematic affinities remain unclear. Another taxon, the presumed genus *Titanodorippe* Blow and Manning, 1996, from the middle Eocene of USA, is a single cheliped propodus that cannot be assigned with confidence to Dorippidae or even Dorippoidea. Among the extant dorippoid genera, only *Ethusa* Roux, 1830 [in Roux, 1828-1830], has its oldest record in the uppermost Eocene (Priabonian, 37.71–33.9 Ma) (Müller and Collins, 1991), while *Dorippe* Weber, 1795, and *Medorippe* Manning and Holthuis, 1991, are known from the lower-middle Miocene (Burdigalian, 20.44–15.97 Ma) (Studer, 1892; Karasawa, 2000). Given the close proximity in age between the occurrences of *Bartethusa* and *Ethusa* from the middle and late Eocene, respectively, the most recent common ancestor of crown Dorippoidea and all of its descendants must have originated during the early Paleogene or even the Cretaceous, while the presence of several potential stem dorippoids in the Early and Late Cretaceous worldwide support a late Mesozoic origin for the superfamily.

## 27. Brachyura: Eubrachyura: Leucosioidea: Leucosiidae (crown)

### Fossil specimen

*Typilobus alponensis* Beschin, De Angeli, and Zorzin, 2009. Museo di Storia Naturale di Verona. Holotype IGVR78539, a complete dorsal carapace in volume (Fig. 5B).

### Phylogenetic justification

The overall oval and domed carapace about as long as wide, with convex anterolateral and posterolateral margins, a narrow posterior margin, the small and round orbits, and the presence of a projected bilobed front/rostrum that is mesially depressed, are typical features of Leucosiidae, an extant family nested within Leucosioidea (Beschin et al., 2009).

*Minimum age*. 47.8 Ma.

*Tip maximum age.* 56.0 Ma.

*Soft maximum age.* 100.9 Ma.

### Age justification

The holotype and sole specimen of *Typilobus alponensis* comes from lower Eocene (Ypresian, 56.0–47.8 Ma) rocks cropping out in the valley of Alpone, near Monte Serea, San Giovanni Ilarione, Verona, Italy (Beschin et al., 2009).

Soft maximum age as for calibration 8.

### Discussion

*Typilobus* Stoliczka, 1871, is an extinct genus with presumably over a dozen species ranging in age from early Eocene (Ypresian) to Miocene and widely distributed across Europe, the UK, North Africa, Asia, and the Indo-Pacific (Karasawa, 1998; Beschin et al., 2009; Artal and Hyžný, 2016; Karasawa et al., 2019). However, several of these species are considerably different from the type species of the genus, *T. granulosus* Stoliczka, 1871, from the Early Miocene of Pakistan, casting doubts on the monophyletic nature of the genus as currently envisioned. Regardless of the generic affinities of *Typilobus alponensis*, this species fits well within the overall body plan seen across crown Leucosiidae (see above under *Phylogenetic justification*), hence our species selection for the calibration of the node. Other Eocene species assigned to *Typilobus* that might be related to *T. alponensis* and thus belong to the crown Leucosiidae are *T. belli* Quayle and Collins, 1981, from the Bartonian of the UK, *T. prevostianus* (Desmarest, 1822, as Leucosia), from the Lutetian of Paris, *T. semseyanus* Lörenthey 1898, from the Lutetian–Priabonian of Hungary and Italy (Lorenthey and Beurlen, 1929; Beschin et al., 1998), and *T. trispinosus* Lörenthey, 1907, from the Lutetian–Priabonian of Egypt (Beschin et al., 1998; Feldmann et al., 2011a; Artal and Hyžný, 2016).

The type material of the monotypic genus *Zannatoius* Beschin, Busulini & Tessier *in* Beschin et al., 2018, from the upper Eocene of Italy, is somewhat damaged and missing the fronto-orbital margin, but overall it conforms with the leucosiid body plan. Among the extant leucosiid genera with known fossil record, *Ebalia* Leach, 1817, has its earliest putative occurrence in the upper Eocene (Priabonian, 37.71–33.9 Ma) of Germany (Förster and Mundlos, 1982; Karasawa et al., 2019). As such, the presence of crown Leucosiidae in the early and late Eocene suggests that the family must have originated at least during the early Paleogene.

## 28. Brachyura: Eubrachyura: Goneplacoidea: Euryplacidae (crown)

### Fossil specimens

*Chirinocarcinus wichmanni* (Feldmann, Casadío, Chirino-Galvez, and Aguirre-Urreta, 1995) (as ?*Glyphithyreus*). Geology Collections at National University of La Pampa, Santa Rosa, La Pampa, Argentina. Holotype GHUNLPam 7015, a nearly complete dorsal carapace (Fig. 5C).

### Phylogenetic justification

*Chirinocarcinus wichmanni* has been included within Euryplacidae based on the relatively broad fronto-orbital margin and large orbits, the well-defined supraorbital angle, and short anterolateral margins (Feldmann et al., 1995; Karasawa and Kato, 2003; Castro and Ng, 2010).

*Minimum age*. 61.6 Ma.

*Tip maximum age.* 66.0 Ma.

*Soft maximum age.* 100.9 Ma.

### Age justification

The type material of *Chirinocarcinus wichmanni* was collected from lowermost Paleocene (lower Danian) rocks of the Roca Formation at the type section, located to the north of General Roca, Rio Negro, Neuquén Province, Argentina (Feldmann et al., 1995). The Roca Formation has been dated as uppermost Cretaceous (Maastrichtian) to lower Paleocene (Danian), based on foraminifera, calcareous nannoplankton, and palynomorphs (del Rio et al., 2011 and references therein). The crab-bearing intervals of the Roca Formation are comprised within the calcareous nannofosil zones NP3 and NP4, with *Chiasmolithus danicus*, *Neochiastozygus modestus*, *Hornibrookina edwardsii*, *H. teuriensis*, and *Toweius africanus* from NP3, indicative of the lower Danian (del Rio et al., 2011).

Soft maximum age as for calibration 8.

### Discussion

Goneplacoidea, as traditionally envisioned, is among the most anatomically disparate and problematic brachyuran groups in terms of the clarity about their internal phylogenetic affinities, and thus their likely polyphyletic relationships with other eubrachyuran groups (Ng et al., 2008; Castro et al., 2010). The extant family Euryplacidae, however, seems to be monophyletic (Castro and Ng, 2010) and nested within the crown Goneplacoidea (Karasawa and Kato, 2003).

Among the euryplacid genera, the monotypic *Chirinocarcinus* Karasawa and Schweitzer, 2004, is the only taxon known from Paleocene fossils. Five other monotypic genera currently assigned to Euryplacidae are known from the lower Eocene (Ypresian) of Italy and the middle Eocene (Lutetian) of Spain (Karasawa and Kato, 2003; Beschin and De Angeli, 2011; Beschin et al., 2016a), as it is the genus *Corallicarcinus* Müller and Collins, 1991, from Eocene fossils from Hungary and Italy (Müller and Collins, 1991; Karasawa and Kato, 2003; Beschin et al., 2018).

Similarly, the genus *Orbitoplax* Tucker and Feldmann, 1990, also referable to Euryplacidae, is known from four Eocene species occurring in Mexico and USA (e.g., Rathbun, 1926; Tucker and Feldmann, 1990; Schweitzer, 2000; Vega et al., 2001).

Among the extant euryplacid genera, *Eucrate* De Haan, 1835, has its oldest putative record in the Oligocene, as represented by a fairly complete ventral carapace with chelipeds of *E. martini* Rathbun, 1926 from USA, and *E. puliensis* Hu & Tao, 1996, from Taiwan. ?*Euryplax culebrensis* Rathbun, 1918 [1919], from the Panama Canal Zone, is of lower Miocene age and not Oligocene, just like the other fossil decapods from the Culebra Formation in the Panama Canal, as initially assumed by Rathbun and repeated by several authors (see in Luque et al., 2017 and comments therein).

The number of extinct and extant euryplacids known from the Paleogene, and the proximity of *C. wichmanni* to the Cretaceous/Paleogene boundary, indicates that Euryplacidae must have a pre-Cenozoic origin, most likely in the Cretaceous.

### 29. Brachyura: Eubrachyura: Eriphioidea: Eriphiidae (crown)

#### Fossil specimens

*Eriphia verrucosa* (Forskal, 1775), in Betancort et al., 2014. Repository indet. Specimen number indet. Single left cheliped fragment with palm and pollex (Fig. 5 D,E).

#### Phylogenetic justification

The type species of *Eriphia* Latreille, 1817, is *E. verrucosa* (Forskal, 1775) [*E. spinifrons* (Herbst, 1782)], by subsequent designation (see in Koh and Ng, 2008; Ng et al., 2008). Although Eriphioidea as previously envisioned is shown to be polyphyletic (see Wolfe et al., 2022), *Eriphia* is the type genus of the family Eriphiidae and thus the superfamily Eriphioidea. As such, it is reliably placed within the crown group.

*Minimum age*. 4.1 Ma.

*Tip maximum age.* 9.3 Ma.

*Soft maximum age.* 100.9 Ma.

#### Age justification

The specimen of *E. verrucosa* reported by Betancort et al., 2014 comes from rocks of the Papagayo cliff, Lanzarote, Canary Islands, Spain, dated as upper Miocene (Messinian?) to lower Pliocene from K/Ar radiometric dating, ranging between 9.3 and 4.1 Ma (see in Betancort et al., 2014, and references therein).

Soft maximum age as for calibration 8.

#### Discussion

The family Eriphiidae, as currently recognized, includes only the extant genera *Eriphia* Latreille, 1817, and *Eriphides* Rathbun, 1897 (Ng et al., 2008; De Grave et al., 2009). While *Eriphides* is monotypic, Eriphia is represented by eight extant and two fossil species (Koh and Ng, 2008; Schweitzer et al., 2010). The extinct *Eriphia cocchii* Ristori, 1886, is known from well-preserved specimens in volume as well as from isolated dactyli from the lower Pliocene (Zanclean– Piacentian, ∼5,33–3.6 Ma) of Italy (Ristori, 1886; De Angeli et al., 2009) (Fig. 5F). Based solely on an isolated propodus, Collins and Donovan, 1997, report a presumably new species, *Eriphia xaymacaensis* Collins and Donovan, 1997, from the upper Pleistocene of Jamaica (Luque et al., 2017). The latter species, however, might be a junior synonym of *Eriphia gonagra* Fabricius, 1781, an extant species. Along these lines, among the extant species with putative fossils, Varola (1981) reported isolated dactyli of the extant species *E. verrucosa* from the mid-Pliocene of Italy, and Luque et al., 2018, reported a cheliped of *Eriphia* cf. *squamata* Stimpson, 1859, from the Quaternary of Panama.

As we show here, the fossil record of Eriphiidae is meager and restricted to the uppermost Neogene and Quaternary, hence why we opt to use a calibration fossil with uncertain specimen data. As such, the most recent common ancestor of extant eriphiids and all of its descendants must have originated at least in the early Neogene or, most likely, in the Palaeogene.

### 30. Brachyura: Eubrachyura: Trapezioidea: Trapeziidae (crown)

#### Fossil specimen

*Archaeotetra lessinea* De Angeli and Ceccon, 2013. Museo Civico “D. Dal Lago” of Valdagno (Vicenza) (MCV). Holotype MCV 12/05-I.G.360311, a complete dorsal carapace (Fig. 5G).

#### Phylogenetic justification

Phylogenetically, Trapeziidae Miers, 1886 seems to be a monophyletic family nested within Trapezioidea s.l. (Castro et al., 2004; Lai et al., 2009; Wolfe et al., 2022), but the monophyly of Trapezioidea as a whole, containing Tetrallidae Castro, Ng, and Ahyong, 2004, has been questioned (see Wolfe et al., 2022). Despite this, the distinctive carapace outline of *Archaeotetra* Schweitzer, 2005, and thus of *Archaeotetra lessinea*, with a wide and slightly bilobed front, the fronto-orbital margin being about 80–90% as wide as the carapace maximum with, the orbits placed anterolaterally, the smooth dorsal carapace, and the seemingly spineless, short, and nearly parallel anterolateral margins, conforms with the body form typical of genera and species in the family Trapeziidae (Schweitzer, 2005; Karasawa and Schweitzer, 2006; De Angeli and Ceccon, 2013; Poore and Ahyong, 2023).

*Minimum age*. 47.8 Ma.

*Tip maximum age.* 53.3 Ma.

*Soft maximum age.* 100.9 Ma.

#### Age justification

The material studied comes from micritic limestones with abundant nullipore coralline algae and fragments of invertebrates, cropping out in the vicinity of Monte Magré, eastern margin of Lessini Mounts, Italy. Age as for calibration 21.

Soft maximum age as for calibration 8.

#### Discussion

Among the few trapeziid crab fossils known, at least four species besides *Archaeotetra lessinea* have been recovered from Eocene deposits. *Archaeotetra inornata* Schweitzer, 2005, is known from the lower to middle Eocene of Mexico, *Paratetralia convexa* Beschin, Busulini, De Angeli, and Tessier, 2007, from the lower Eocene of Italy, and *Eomaldivia pannonica* and *E. trispinosa* Müller and Collins, 1991, from the upper Eocene of Hungary. This indicates that the most recent common ancestor of trapeziid crabs and all of its descendants must have a pre-Eocene origin (Wolfe et al., 2022).

### 31. Brachyura: Eubrachyura: Eriphioidea: Oziidae (crown)

#### Fossil specimen

*Ozius collinsi* Karasawa, 1992. Mizunami Fossil Museum (Yamanouchi, Mizunami, Japan). Holotype MFM39001, a complete dorsal carapace (Fig. 5H).

#### Phylogenetic justification

The extant genus *Ozius* Milne Edwards, 1834 [in H. Milne Edwards, 1834-1840] is the type genus of the family Oziidae. The overall anatomy of *Ozius collinsi* fits well the diagnosis of the genus, and thus can be considered within the crown Oziidae s.s.

***Minimum age***. 13.82 Ma.

*Tip maximum age.* 15.97 Ma.

*Soft maximum age.* 100.9 Ma.

#### Age justification

The type material of *Ozius collinsi* comes from mudstones of the Yoshino Formation, Katsuta Group, Locality KTT13, in Niida, Tsuyama City, Okayama Prefecture, Japan (Karasawa, 1992). The Yoshino Formation is considered as uppermost lower Miocene to middle Miocene (Burdigalian to Langhian), with some estimated ages of 17.9± 2.1 Ma and 18.5–16 Ma based on the fission track method and palynomorphs (Mori and Yamanoi, 2003; Suzuki et al., 2003, in Ando et al., 2016), and on mollusc assemblages and planktonic foraminifera biostratigraphic ages, zones N8 to N10 (Blow, 1969; Yoshimoto, 1979, in Karasawa, 1992 and Taguchi, 2002), which correspond to the global Burdigalian. Other authors have restricted the age of the oldest fossil Oziidae from Japan to the Langhian (Schweitzer et al., 2021b).

Soft maximum age as for calibration 8.

#### Discussion

In some recent mitogenomic studies, oziid taxa such as *Epixathus frontalis* have been recovered outside the traditional Eriphioidea, highlighting the polyphyletic nature of the superfamily as traditionally envisioned (Tan et al., 2018; Lü et al., 2022), and further corroborated in Wolfe et al. (2022). Moreover, previous groupings of genera within the family Oziidae have been shown to be non-monophyletic (Lai et al., 2014), further adding to the phylogenetic issues highlighted above.

This also adds to the problem that several extinct forms that might superficially resemble oziids might not necessarily belong to Oziidae s.s., which, together with their incompleteness, preclude a confident systematic placement. As such, due to the considerably reliable placement of *O. collinsi* within the genus *Ozius*, and due to its early-mid Miocene age, we consider it as the best calibration point for the family currently available.

### 32. Brachyura: Eubrachyura: Pilumnoidea: Pilumnidae (crown)

#### Fossil specimen

*Glabropilumnus trispinosus* Beschin, Busulini, and Tessier, *in* Beschin et al., 2016a. Museo di Storia naturale di Verona. Holotype VR 94508, a complete dorsal carapace in volume (Fig. 5I).

#### Phylogenetic justification

The overall carapace shape of *G. trispinosus*, with smooth dorsal surface and lacking dorsal ridges, and the anterolateral margin with 3 spines, being the posterior one the smallest of all, match well the diagnosis of *Glabropilumnus* (Poore and Ahyong, 2023). *Glabropilumnus* Balss, 1932, is an extant genus represented by six extant species (Ng et al., 2008), and the only genus in the Family Pilumnidae Samouelle, 1819, and the subfamily Pilumninae Samouelle, 1819, with a fossil record extending into the Eocene (see below), hence our taxon selection for calibration.

*Minimum age*. 47.8 Ma.

*Tip maximum age.* 56.0 Ma.

*Soft maximum age.* 100.9 Ma.

#### Age justification

*Glabropilumnus trispinosus* is known from its type locality in the area of Rama di Bolca (Verona), NE Italy, in calcareous rocks of the ‘Calcari Nummulitici’ that have been ascribed to the lower Eocene (Ypresian) based of the presence of *Nummulites partschi*, *Alveolina cremae*, *A. croatica*, *A. decastroi*, *A*. cf. *dainellii*, and *A. distefanoi* (Beschin et al., 2016a, and references therein).

Soft maximum age as for calibration 8.

#### Discussion

Pilumnid genera and species show a considerable anatomical disparity. Although previous molecular and mitogenomic phylogenetic studies have suggest that Pilumnidae, in the broad sense, is monophyletic (Lai et al., 2014; Duan et al., 2022), the monophyly of Pilumnidae as currently envisioned is questionable (Wolfe et al., 2022). Despite this, *Glabropilumnus* can be referred to the subfamily Pilumninae in the strict sense, and thus to the crown Pilumnidae (Poore and Ahyong, 2023).

There are several extinct genera of Eocene crabs from Europe and USA that have been tentatively assigned to Pilumnidae and, specifically, the subfamily Pilumninae. However, the only extant pilumnid genus with Eocene representatives known is *Glabropilumnus* Balss, 1932. Besides *G. trispinosus, G. bizzarinii* Beschin, Busulini, and Tessier, in Beschin et al., 2019, is known from the upper Eocene (Priabonian) of Italy, together with *G.* cf. *granulatus* De Angeli, Garassino & Ceccon, 2010b, which was previously known solely from the lower Oligocene of Italy (Beschin et al., 2019). As such, the most recent common ancestor of Pilumnidae and all of its descendants must be pre-Eocene.

### 33. Brachyura: Eubrachyura: Goneplacoidea: Goneplacidae (crown)

#### Fossil specimens

*Carcinoplax temikoensis* Feldmann and Maxwell, 1990. New Zealand Geological Survey, Lower Hutt, New Zealand. Holotype AR 1943, a complete dorsal carapace and a cheliped (Fig. 5J).

#### Phylogenetic justification

*Carcinoplax temikoensis* can be assigned to *Carcinoplax* Milne Edwards, 1852, based on the transversely rectangular carapace, the forward directed, narrow orbits, and the presence of three spines on the anterolateral margin, including the one forming the outer orbital angle (Feldmann and Maxwell, 1990; Ng and Castro, 2020). *Carcinoplax* is nested within Goneplacidae (Karasawa and Kato, 2003).

*Minimum age*. 34.6 Ma.

*Tip maximum age.* 39.1 Ma.

*Soft maximum age.* 100.9 Ma.

#### Age justification

The type material of *Carcinoplax temikoensis* was collected from the lower part (unit 2) of the Island Sandstone at Te Miko, locality K30/D7A, exposed along the coast between Perpendicular Point and Punakaiki, north of Greymouth, Westland, New Zealand, ascribed to the upper Eocene (Kaiatan to Runangan) based on foraminiferal assemblages (Feldmann and Maxwell, 1990). The Kaiatan (39.1–36.7 Ma) and Runangan (36.7–34.6 Ma) stages in New Zealand roughly correspond to the global uppermost Bartonian (41.2–37.71 Ma) and Priabonian (37.71–33.9 Ma) ages (Raine et al., 2015). Given the unclear exact age of the layers bearing the type material of *C. temikoensis*, for the time being, we bracket its tip maximum and minimum ages between the base of the Kaiatan and the top of the Runangan (i.e., 39.1–34.6 Ma).

Soft maximum age as for calibration 8.

#### Discussion

Currently, there are 25 species referred to *Carcinoplax* (Ng and Castro, 2020). As previously discussed for Euryplacidae (see above), the superfamily Goneplacoidea is an anatomically disparate and problematic brachyuran groups in terms of the clarity about its internal phylogenetic affinities (Ng et al., 2008; Castro et al., 2010). Despite this, *Carcinoplax* fits well within Goneplacidae and thus belongs to the crown Goneplacoidea. There are over a dozen species of *Carcinoplax* known from fossils, from which only *C. temikoensis* occur in the Eocene. Extinct, monotypic genera known from the Eocene and presumably with goneplacid affinities include *Amydrocarcinus* Schweitzer, Feldmann, González-Barba, and Vega, 2002, from the Bartonian of Mexico, and *Gonioplacoides* Quayle and Collins, 2012, from the upper Eocene of UK. The presence of extinct and extant goneplacid genera in the Eocene indicate that the most recent common ancestor of Goneplacidae and all of its descendants must be at least early Eocene in age, but, as previously seen with most other families already discussed, its origins might lie in the Paleocene (Wolfe et al., 2022).

### 34. Brachyura: Eubrachyura: Xanthoidea: Pseudorhombilidae (crown)

#### Fossil specimens

*Pseudorhombila patagonica* Glaessner, 1933. Collections of the British Museum. Holotype In. 28031, a complete dorsal carapace (Fig. 5K).

#### Phylogenetic justification

The genus *Pseudorhombila* H. Milne Edwards, 1834 [in 1834-1840], has been recovered nested within a monophyletic Xanthoidea, but its familial placement has been included either within Pseudorhombilidae Alcock, 1900 (e.g., Thoma et al., 2014), or within the subfamily Pseudorhombilinae under the family Panopeidae Ortmann, 1893, and besides other subfamilies such as Panopeinae, Melybiinae, and an unnamed subfamily (e.g., Mendoza et al., 2022). In our study (Wolfe et al., 2022), *Pseudorhombila* was recovered in a clade formed by several genera typically ascribed to Pseudorhombilidae, which in turn was recovered as sister to a clade formed by several genera ascribed to Panopeidae. As such, here we consider Pseudorhombilidae as a discrete family.

***Minimum age***. 5.333 Ma.

*Tip maximum age.* 23.03 Ma.

*Soft maximum age.* 100.9 Ma.

#### Age justification

The type specimen of *Pseudorhombila patagonica* was collected from the so-called Patagonian Beds, in the Santa Cruz Province, Argentina, which have been considered Miocene in age (Glaessner, 1933, 1969). Unfortunately, it comes from an old collection, and its precise lithostratigraphic and chronostratigraphic context is unclear, hence the broad minimum and soft maximum age bracket.

Soft maximum age as for calibration 8.

#### Discussion

Pseudorhombilidae Alcock, 1900, is a small family of xanthoid crabs with eight extant genera and over a dozen species known to date (Ng et al., 2008; De Grave et al., 2009). In our study (Wolfe et al., 2022), species within Pseudorhombilidae form a monophyleic group sister to Panopeidae. To date, the only known fossil referable to the family is *P. patagonica*, a member of the extant genus *Pseudorhombila* H. Milne Edwards, 1834 [in 1834-1840], therefore it is our selection of fossil for calibration.

### 35. Brachyura: Eubrachyura: Xanthoidea: Panopeidae (crown)

#### Fossil specimens

*Panopeus incisus* Beschin, Busulini, De Angeli, and Tessier, 2007. Museo di Archeologia e Scienze Naturali “G. Zannato” di Montecchio Maggiore, Vicenza, northern Italy. Holotype MCZ 2009, a nearly complete dorsal carapace (Fig. 5L).

#### Phylogenetic justification

*Panopeus incisus* was ascribed to *Panopeus* H. Milne Edwards, 1834 [in 1834-1840], based on the wide, subhexagonal carapace, the broad and mesially notched front, the sub-ovate orbits bearing two orbital fissures, and the well-developed anterolateral margin with four spines excluding the outer orbital spine (Schweitzer, 2000; Beschin et al., 2007). *Panopeus* is an extant genus, the type genus of the family Panopeidae Ortmann, 1893, and together with other panopeid species forms a clade nested within Xanthoidea (Jennings et al., 2021; Mendoza et al., 2022Thoma et al., 2014; Wolfe et al., 2022).

*Minimum age*. 47.8 Ma.

*Tip maximum age.* 56.0 Ma.

*Soft maximum age.* 100.9 Ma.

#### Age justification

The type material of *Panopeus incisus* comes from lower Eocene rocks cropping out at the Contrada Gecchelina of the Monte di Malo in Vicenza, northern Italy (Beschin et al., 2007, see also Beschin et al., 2016a). Age as for calibration 13.

Soft maximum age as for calibration 8.

#### Discussion

Panopeidae is a speciose group of xanthoid crabs with an early fossil record extending back into the Paleogene Period. Among the extant panopeid genera, only *Panopeus*, *Lophopanopeus* Rathbun, 1898, and *Metopocarcinus* Stimpson, 1860, have putative occurrences in the Eocene (Beschin et al., 2016b; Pasini et al., 2019), with *Panopeus* being represented by multiple species (e.g., Schweitzer, 2000; Vega et al., 2008; Gatt and De Angeli, 2010; Beschin et al., 2013; Beschin et al., 2016a; Beschin et al., 2018, and references therein). While there are numerous extinct presumed panopeid genera with Eocene representatives, only *Pakicarcinus* Schweitzer, Feldmann and Gingerich, 2004, and *Glyphithyreus* Reuss, 1859, have pre-Eocene (upper? Paleocene) occurrences (Collins and Morris, 1978; Charbonnier et al., 2013; Schweitzer et al., 2016b). The presence of numerous species of extinct and extant panopeid genera in the Eocene and apparently the late Paleocene, suggest that the origins of the family are rooted into the early Paleogene.

### 36. Brachyura: Eubrachyura: Xanthoidea: Xanthidae (crown)

#### Fossil specimens

*Phlyctenodes edwardsi* Beschin, Busulini, Tessier, and Zorzin, 2016a. Museo di Storia naturale di Verona. Holotype VR 94284, a complete dorsal carapace (Fig. 5M).

#### Phylogenetic justification

The genus *Phlyctenodes* Milne-Edwards, 1862a is ascribed to Xanthidae, specifically the subfamily Actaeinae Alcock, 1898, based on the ovate and wide carapace with poorly defined dorsal regions, the large and rimmed sub-circular orbits lacking orbital fissures, the anterolateral margins moderately convex and bearing several small tubercles or spines, the smooth posterolateral margins nearly straight to weakly concave, the overall fronto-orbital margin configuration, a front that is wide and bearing four or more tubercles, and the overall outline of the frontal and anterolateral margins forming a wide arch (Busulini et al., 2006; Beschin et al., 2016a; Schweitzer et al., 2021d).The monophyletic status of the family Xanthidae MacLeay, 1838b, has been corroborated by several molecular and total evidence phylogenetic studies (e.g., Lai et al., 2011; Tsang et al., 2014; Mendoza et al., 2022; Wolfe et al., 2022).

*Minimum age*. 47.8 Ma.

*Tip maximum age.* 56.0 Ma.

*Soft maximum age.* 100.9 Ma.

#### Age justification

*Phlyctenodes edwardsi* is known from its type locality in the area of Rama di Bolca (Verona), NE Italy, in calcareous rocks of the ‘Calcari Nummulitici’ that have been ascribed to the lower Eocene (Ypresian) (Beschin et al., 2016a). Age as for calibration 32.

Soft maximum age as for calibration 8.

#### Discussion

*Phlyctenodes* is an extinct genus with most of its species occurring in the Eocene of Europe (Busulini et al., 2006). We chose *P. edwardsi* as our calibration point given its Ypresian (56.0–47.8 Ma) age, which is similar to the age assigned to *P. multituberculatus* Beschin, Busulini, De Angeli, and Tessier, 2007, and *P. nicolisi* Bittner, 1884, from the Ypresian of Italy. *Phlyctenodes tuberculosus* Milne-Edwards, 1862a, from the middle–upper Eocene (41.2–47.8 Ma) of France and Italy, was used by Tsang et al. (2014), Wolfe et al. (2019), and Mendoza et al. (2022) as their calibration points. Beschin et al. (2016a) reported this species also from the Ypresian of Italy, in the Bolca region from were *P. edwardsi* originated. As such, given the numerous species of *Phlyctenodes* now know from the Ypresian, any of those species would represent equally parsimonious choices of calibration points, and all would indicate that the most recent common ancestor of xanthid crabs and all of its descendants must have originated in the pre-Eocene.

## ACKNOWLEDGEMENTS

We thank Matúš Hyžný, Àlex Ossó, Barry van Bakel, Fernando Ari Ferratges, Carrie Schweitzer, and Orangel Aguilera, for facilitating us bibliographic references and for valuable discussion about the systematic affinities and potential ages of some of taxa here investigated. Special thanks to Pierre Olivier Antoine (Université de Montpellier, France), Pedro Artal (Museo Geológico del Seminario de Barcelona, Spain), Barry van Bakel (Oertijdmuseum De Groene Poort, Boxten, the Netherlands), Juan Francisco Betancort (Universidad de Las Palmas de Gran Canaria, Spain), Lisa Boucher and Ann Molienux (UT Austin, USA), Alessandra Busulini (Museo di Storia Naturale, Venezia, Italy), Antonio De Angeli (Museo Civico “G. Zannato”, Montecchio Maggiore, Vicenza, Italy), Alfréd Dulai (Hungarian Natural History Museum, Hungary), Rodney Feldmann and Carrie Schweitzer (Kent State University, USA), Fernando Ari Ferratges (Universidad de Zaragoza, Spain), Richard Howard (NHM London, UK), Matus Hyzny (Comenius University, Slovakia), Hiroaki Karasawa (Mizunami Fossil Museum, Japan), Adiël Klompmaker (Alabama), Àlex Ossó (Tarragona, Catalonia), and Francisco Vega (UNAM, Mexico), for their kind help providing several of the images of the fossil specimens here illustrated. We thank the National Science Foundation (NSF), DEB grants #1856667 and #1856679 (USA) for supporting this work. This is contribution #X from the Coastlines and Oceans Division of the Institute of Environment at Florida International University.

## TABLE CAPTIONS

**Table 1.** Summary list of 40 fossil meiurans and 36 vetted fossils for molecular divergence time estimations of brachyuran crabs (one anomuran outgroup, 35 brachyuran ingroups). Ages in millions of years (Ma).

